# Genome-wide molecular effects of the neuropsychiatric 16p11 CNVs in an iPSC-to-iN neuronal model

**DOI:** 10.1101/2020.02.09.940965

**Authors:** Thomas R. Ward, Xianglong Zhang, Louis C. Leung, Bo Zhou, Kristin Muench, Julien G. Roth, Arineh Khechaduri, Melanie J. Plastini, Carol Charlton, Reenal Pattni, Steve Ho, Marcus Ho, Yiling Huang, Joachim F. Hallmayer, Phillippe Mourrain, Theo D. Palmer, Alexander E. Urban

**Affiliations:** Department of Psychiatry and Behavioral Sciences, Stanford University School of Medicine, Stanford, CA 94305, USA; Department of Genetics, Stanford University School of Medicine, Stanford, California 94305, USA; Stanford Center for Sleep Sciences and Medicine, Stanford University School of Medicine, Stanford, CA 94305, USA; Department of Neurosurgery, Stanford University School of Medicine, Stanford, CA 94305, USA

**Keywords:** 16p11.2, genomic deletion, genomics duplication, PCSK9, gene expression, DNA methylation

## Abstract

Copy number variants (CNVs), either deletions or duplications, at the 16p11.2 locus in the human genome are known to increase the risk for autism spectrum disorders (ASD), schizophrenia, and for several other developmental conditions. Here, we investigate the global effects on gene expression and DNA methylation using a 16p11.2 CNV patient-derived induced pluripotent stem cell (iPSC) to induced neuron (iN) cell model system. This approach revealed genome-wide and cell-type specific alterations to both gene expression and DNA methylation patterns and also yielded specific leads on genes potentially contributing to some of the known 16p11.2 patient phenotypes. PCSK9 is identified as a possible contributing factor to the symptoms seen in carriers of the 16p11.2 CNVs. The protocadherin (PCDH) gene family is found to have altered DNA methylation patterns in the CNV patient samples. The iPSC lines used for this study are available through a repository as a resource for research into the molecular etiology of the clinical phenotypes of 16p11.2 CNVs and into that of neuropsychiatric and neurodevelopmental disorders in general.

## Introduction

Neuropsychiatric disorders have a high prevalence in humans, ranging from ∼.57% for Schizophrenia to 19.1% for any Anxiety Disorder in the US based on reports from the National Institute of Mental Health. All of these disorders are estimated to have high levels of heritability. While the specific underlying molecular mechanisms are still mostly unknown, multiple regions of the genome as well as increasing numbers of candidate genes of have been found to be associated with these disorders. Several high-confidence candidate loci now exist and those with the highest penetrance are large copy number variants (CNVs) (Stankiewicz and Lupski, 2010; Sullivan et al., 2012).

CNVs are large deletions or duplications in the human genome that are at least 1 kb in size but that often range into the hundreds of thousands to millions of basepairs. Disease relevant CNVs often occur *de novo* (Levy et al., 2011), possibly through non allelic homologous recombination (NAHR) events due to the deleted or duplicated regions being flanked by segmental duplication regions (SegDups), also known as low copy repeats (LCRs). Many small-to-medium-sized CNVs are common in the population and have no known phenotypic effects (Iafrate et al., 2004) but there are around 70 to 120 CNV regions that are associated with genomic disorders (Girirajan et al., 2012). These disease associated CNVs, such as for example large CNVs on chromosomes 22q11.2, 1q21.2, and 15q13.3, are a prominent genetic cause in 5-15% of individuals suffering from intellectual deficit, developmental delay (DD), autism spectrum disorders (ASD), congenital malformation (CM), among others (Girirajan et al., 2013).

Two large CNVs with the strongest association to neuropsychiatric and generally neurological disorders are the 16p11.2 BP4-BP5 deletion CNV (MIM 611913) and the duplication CNV (MIM 614671) in the same locus, which are known to be present in 1% of Autism spectrum disorder (ASD) patients. Furthermore the duplication CNV in this locus increases the risk for developing Schizophrenia by 14-fold (Deshpande and Weiss, 2018; Miller et al., 2015). The 600kb CNV with start and endpoints within the same SegDups can occur as either a deletion or a duplication and is present in at least 3/10,000 individuals (Weiss et al., 2008; Bijlsma et al., 2009). Along with the neuropsychiatric disorders, the 16p11.2 CNV has been associated with several other developmental phenotypes including speech and language delay, seizures, intellectual disability (ID), developmental delay (DD), developmental coordination disorder (DCD), anxiety, and ADHD (Ghebranious et al., 2007; Kumar et al., 2008; Fernandez et al., 2010; Rosenfeld et al., 2010; Shinawi et al., 2010; Zufferey et al., 2012; Hanson et al., 2015; Green Snyder et al., 2016; Steinman et al., 2016; Bernier et al., 2017). Additionally, the deletion and duplication cases are known to often show reciprocal phenotypes for head circumference and body mass index (BMI). Specifically, the deletion patients will generally develop macrocephaly, whereas the duplication patients tend to develop microcephaly (Steinman et al., 2016). This trend shows some variance, with 10% of duplication carriers show microcephaly and 17% of deletion carriers showing macrocephaly (Steinman et al., 2016). For BMI, deletion carriers show an increase in BMI by age 7 resulting in ∼75% of adult carriers being obese. Conversely, duplication carriers are at an eightfold risk of being underweight (Jacquemont et al., 2011).

Previous studies of the molecular and cellular impact of these CNVs have already generated important insights but also have left many fundamental questions unanswered. The problem is particularly challenging given the large degree of phenotypic diversity and variance in the patients, the large size of the genomic region affected and the concurrent large number of genes that are encoded within that region. On top of the variance and noise generated in the data from the region, there is also a lack of human neuronal sample tissue to perform these studies in. Previously, human LCLs derived from patients (Blumenthal et al., 2014; Kusenda et al., 2015; Migliavacca et al., 2015) or mouse model systems (Arbogast et al. 2016, Horev et al. 2011) have been used. LCLs stem from the B-lymphocyte cells, therefore insights gained in that model system have to be further explored in cell types closer to those found in the brain. The 16p11.2 mouse models also have yielded many interesting insights already. There are some limitations for this model since the gene content of the CNV in the mouse genome is not entirely identical to that in the human patients. And also there are some notable discrepancies on the phenotypic level, for example the 16p11.2 deletion mouse model shows reduced brain weight which is the opposite of what is seen in human deletion patients where there is often macrocephaly (Horev 2011, Portmann 2014, Pucilowska 2015).

Recently, additional important work has emerged, namely induced pluripotent stem cells (iPSC) have been used to engineer the 16p11.2 CNV into a control cell line (Tai DJ et al., 2016), and contrasting cellular phenotypes have been described in patient iPSC-derived neurons (Deshpande et al., 2017). Here we perform a study on the alterations to gene expression and DNA methylation while 16p11.2 CNV patient iPS cell lines differentiate into a neuronal cell type. Using an available resource for 16p11.2 CNV patient iPSC cell lines from the Simons Foundation, we were able to evaluate the cellular impact of the 16p11.2 CNV and identify potential key players in the molecular etiology of some of the clinical phenotypes seen in patients.

## Results

### 16p11.2 CNV patient-derived iPSCs form the basis of induced neurons (iNs)

Using the patient iPSC-to-iN model system (Figure 1A), all cell lines were cultured starting at the same time and were differentiated in batches based on the growth rate and confluency of the cells. Low-coverage whole-genome sequencing was performed on the iPSCs and, within the CNV region, showed a decrease in sequencing coverage for the deletion patients and an increase in sequencing coverage for the duplication patients when compared to the control lines (Figure 1B). The presence of the expected copy number changes was further verified through ddPCR, using TaqMan copy number assays to validate the copy number of the 16p11.2 CNV region (Figure S1). While the cells underwent the differentiation process, GFP imaging and staining of neuronal cell markers TUB and MAP were performed, which confirmed that the differentiated cells were of a neuronal cell type (Figure 1C). WGCNA pathway analysis of the RNA-Seq data was carried out on the list of genes upregulated in the iNs compared to the iPSCs and multiple neuronal pathways were shown to be upregulated (Figure S2).

**Figure 1:**
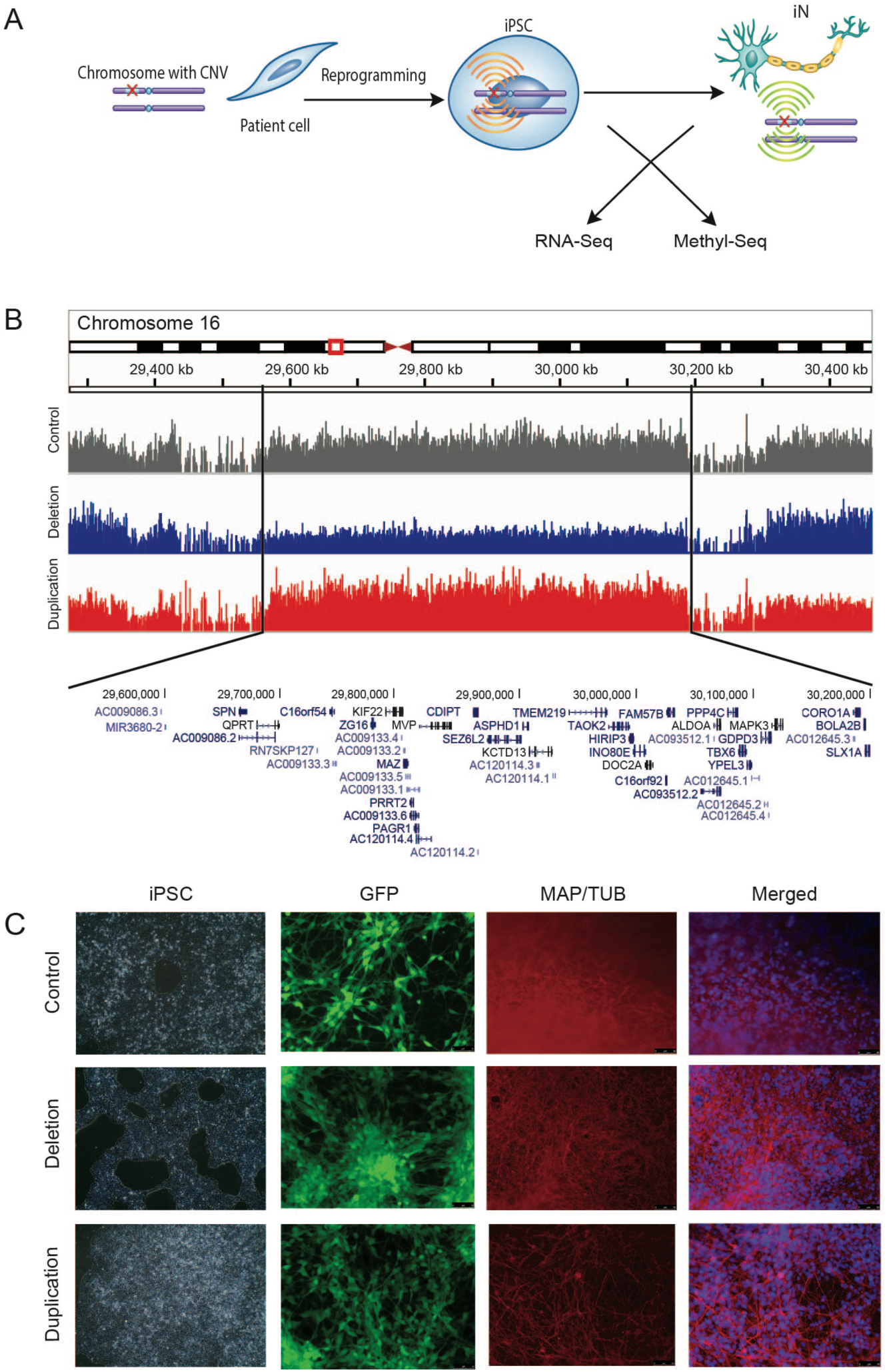
A model system to investigate the 16p11.2 CNVs. **(A)** A cell-model system to study the effects of the CNV in a disease-relevant cell type while. Patient fibroblast cells are reprogrammed into iPSCs which are then differentiated into induced neurons (iNs). Gene expression and DNA methylation patterns are all analyzed to investigate the molecular effects of harboring the CNVs. **(B)** Low-coverage whole-genome sequencing reads of the 16p11.2 CNV region, shown here for each a single control, deletion, and duplication sample, visualized in IGV (Robinson et al. 2011). The deletion line (blue) has lower sequencing coverage while the duplication line (red) has higher sequencing coverage compared to the control cell line (gray), verifying the presence of the CNV. Vertical lines designate the boundaries of the CNV and the genes encoded in the region are shown. **(C)** Fluorescence images were taken of the iNs to verify the successful integration of the viral vectors in the differentiation protocol. First column are images of the iPSC stage for one of each of the controls, deletions, and duplications. Second column shows actively expressing GFP indicating successful integration of the vectors. The iNs were also stained for neuronal differentiation markers MAP and TUB to ensure completion of the differentiation protocol (third column). iNs were co-stained with DAPI and the images were merged (fourth column).

RNA-Seq and CpG-capture bisulfite sequencing were performed on both the control cell lines and the 16p11.2 deletion and duplication patient derived cell lines at both the iPSC and iN cell stages. This approach allowed us to study the molecular effects of the 16p11.2 CNVs in a disease relevant tissue while modeling the developmental conditions during which the patient phenotypes may begin to arise. For the RNA sequencing analyses, all control and patient lines (with the exception of 1 duplication clonal line) were used, for a total of 15 samples: 4 control, 6 deletion (i.e. 2 clonal lines each from 3 patients), 5 duplication (i.e. 2 clonal lines from 2 patients and 1 clonal line from one more patient). A subset of those lines was used for the DNA-methyl-seq analyses, with 2 controls, 2 deletion patient lines, and 2 duplication patient lines: 6632.4, 726.1, 14758×3 101.7, 14765×2 101.2, 14756×9 201.2, and 14723×10 202.8.

### Gene expression levels altered within the CNV region as expected

RNA-Seq was performed for the patient-derived iPSC and iN cell lines, obtaining approximately 20 million Illumina paired-end reads for each sample. The data was analyzed using two analysis pipelines, Cuffdiff and DESeq2. The Cuffdiff analysis produced the main dataset for this project, with DESeq2 being used to confirm the general findings from Cuffdiff (as well as to generate the plots in Figure 2). The most likely scenario regarding gene expression levels is that the deletion of one copy of a gene will result in a decrease in expression level while its duplication will result in a 50% increase. In line with this expectation, we found that the majority of the genes within the CNV region have a change in gene expression that followed the change in genomic copy number, at both the iPSC and iN stages (Figure 2 A, B). The deletion analyses have a consistent change at both cell stages, −0.55 ± 0.10 at iPSC stage and −0.54 ± 0.06 at iN stage. The duplication analyses were more variable in comparison, 2.15 ± 1.05 at the iPSC stage and 1.94 ± 0.31 at the iN stage.

**Figure 2:**
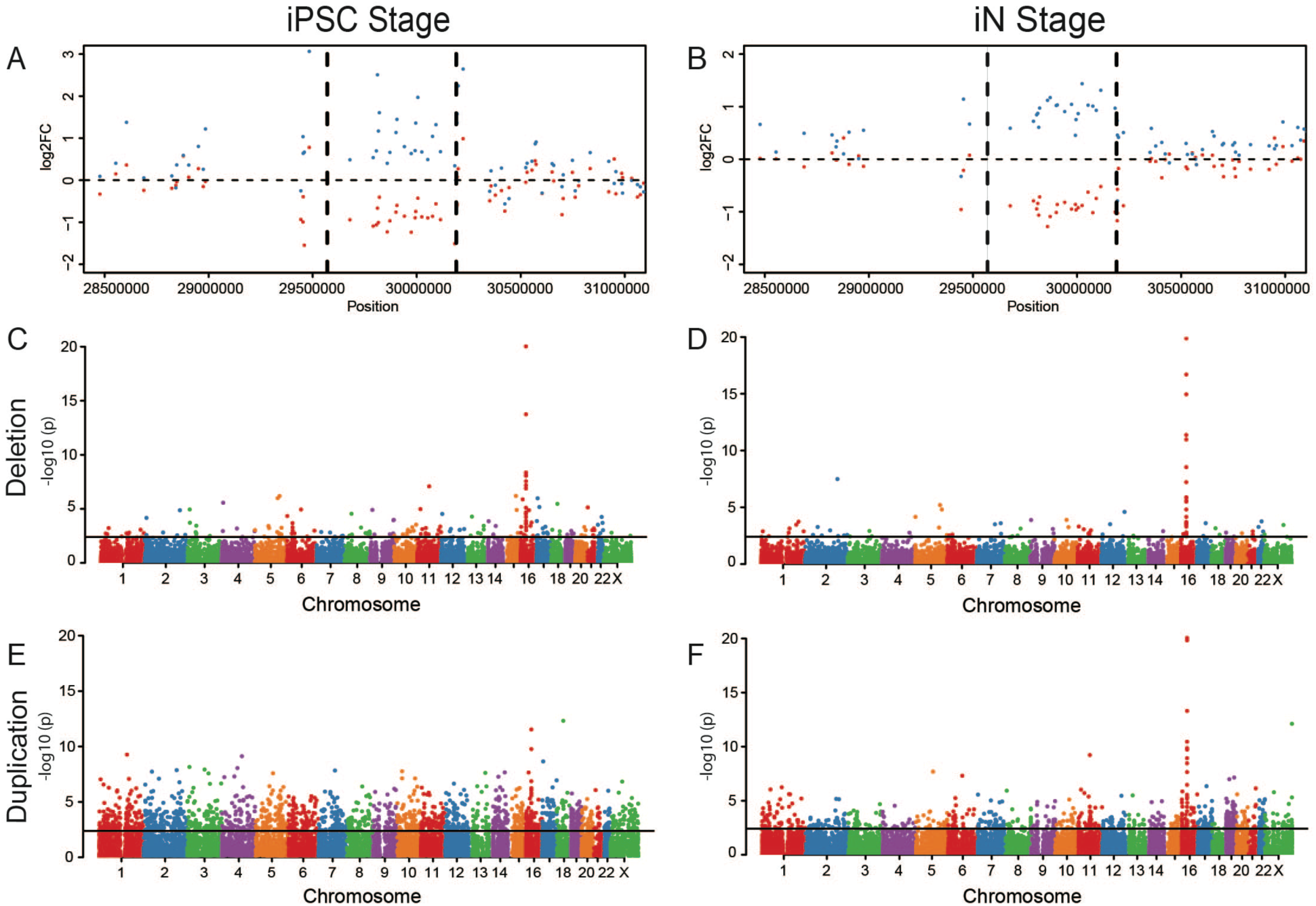
Gene expression is altered within the CNV region and genome-wide. Plots of genes’ log2 fold change in the CNV and flanking regions **(A & B)** and fold change associated –log10(p) across the genome **(C-F)**. **(A & B)** Gene expression within the CNV and the flanking regions, the CNV boundaries are represented by the vertical lines. The deletion analyses showed a decrease in expression (red dots) while the duplication analyses showed an increase in expression (blue dots). Genome-wide, genes reach statistically significant differential expression in the **(C)** iPSC deletion, **(D)** iN deletion, **(E)** iPSC duplication, and **(F)** iN duplication analyses.

### Gene expression is altered genome-wide outside the CNV region

Gene expression alterations were not localized to only the CNV region. Significant differential gene expression was found genome-wide for both CNV types at both cell stages (Figure 2 C-F). At the iPSC stage, iPSC-Del and iPSC-Dup had 16,653 and 16,833 genes with detectable expression, respectively. Of those genes, there were 121 differentially expressed genes (DEGs) for iPSC-Del with 83 of those DEGs being up-regulated and 38 down-regulated. 654 DEGs were in the iPSC-Dup analysis with a split of 341 up-regulated genes and 313 down-regulated genes. At the iN stage, genes with detectable expression decreased moderately in number from the iPSC stage to 16,165 for iN-Del and 16,106 for iN-Dup. The iN-Del analysis had 35 DEGs (29 up-regulated, 6 down-regulated) while 59 DEGs were in the iN-Dup analysis (50 up-regulated, 9 down-regulated). The DEGs from each analysis were also categorized based on genomic location to determine what percentage of these DEGs resided on each chromosome. This analysis revealed multiple chromosomes at both cell stages with a larger percentage of DEGs on them than chromosome 16. At the iPSC stage, the lists of DEGs from the iPSC Del and iPSC Dup analyses combined for a larger percentage of the total DEGs on chromosomes 1, 2, 5, 6, and 19. At the iN stage, this was also the case for chromosomes 1, 11, 19, and 22. Notably, this also does not exclude the DEGs within the CNV region, which contributed significantly to the number of DEGs found on chromosome 16.

There were 65 DEGs overlapping between deletion and duplication lines at the iPSC stage (iPSC-Del + iPSC-Dup) and 9 genes were overlapping at the iN stage (iN-Del + iN-Dup) (Figure 3). At the iPSC stage, PCSK9 was found to be up-regulated in the deletion samples (1.10 log2 fold change) and down-regulated in the duplication samples (−1.16 log2 fold change). At the iN stage, PCSK9 becomes up-regulated (1.72 log2 fold change) in the duplication samples and, while it is not a DEG in the iN Del analysis, it is also up-regulated (1.42 log2 fold change). Gene expression changes were verified using qPCR (Table S1).

**Figure 3:**
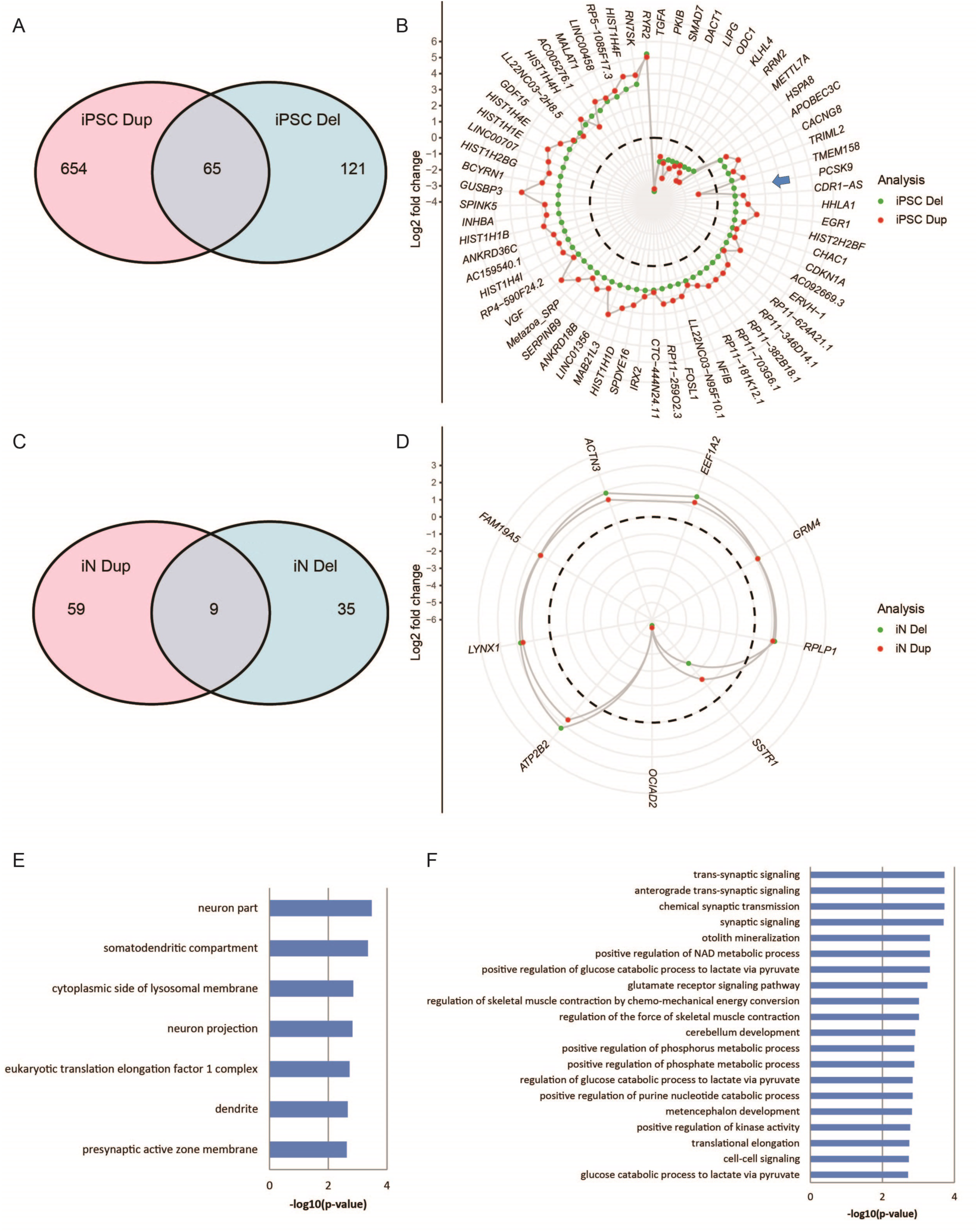
DEGs overlaps between cell stage analyses and functional enrichment analysis. Differentially expressed genes (DEGs) found in the control vs patient sample analyses were compared to find any genes that overlapped at each cell stage. Venn diagrams showing the number of DEGs from each of the four analyses and the number of DEGs that overlap at the **(A)** iPSC stage and the **(C)** iN stage. Log2 fold change was plotted for each DEG from both of the overlapping analyses. Fold change values from the deletion analysis are represented by green and duplication analysis values are represented by red. DEGs overlapping between deletion and duplication patients at the iPSC stage **(B)** and iN stage **(D)**. The dotted line represents zero fold change difference in expression. With the nine DEGs overlapping between the iN analyses, functional enrichment analysis was done using ToppFun which is a part of ToppGene Suite (Chen et al., 2009). **(E)** The output for GO: Cellular Component and the **(F)** top 20 output by p-value for GO:Biological Process are plotted by their corresponding enrichment p-value, converted to –log10(p-value) for the plot.

In order to validate that the fold changes found in the overlapping DEGs, an alternative RNA-Seq pipeline (DESeq2) was used to analyze our data. The four comparison analyses were repeated in DESeq2 and the DEG lists were compiled by selecting the genes with a FDR ≤ .05. Plotting the overlapping DEGs as before, we see a similar pattern of consistent fold changes in each plot with 73 DEGs overlapping at the iPSC stage and 5 DEGs overlapping at the iN stage (Figure S3).

### Overlapping DEGs at the iN stage are relevant for neuronal function

We carried out functional enrichment analysis on the overlapping DEGs at the iN stage using ToppFun, a part of ToppGene Suite (Chen et al., 2009). The analysis yielded several pathways relevant for neuronal function, neuronal processes, and neuronal components, particularly in the GO categories “Biological Process” and “Cellular Component”. The “Cellular Component” output (Figure 3E) showed an enrichment of neuron components such as *neuron part* (p-value 3.319E-4), *somatodendritic compartment* (p-value 4.458E-4), *neuron projection* (p-value 1.496E-3), and *dendrite* (p-value 2.156E-3). The “Biological Process” output (Figure 3F) identified an enrichment of processes related to synapse function such as *chemical synaptic transmission*, *anterograde trans-synaptic signaling*, *trans-synaptic signaling* (p-value 1.895E-4 for previous three), and *synaptic signaling* (p-value 1.994E-4), as well as *cerebellum development* (p-value 1.223E-3).

To ascertain the existence of a group of overlapping DEGs between deletion and duplication tissues in an alternative model system we reanalyzed the RNA-Seq data from the 2014 Blumenthal et al. study in which RNA had been extracted from the cortex of the 16p11.2 deletion and duplication mouse models, respectively, and subjected to RNA-Seq analysis relative to control animals. RNA-Seq data of 4 deletion mice had been compared to 4 control mice and RNA-Seq data of 4 duplication mice had been compared to a second group of 4 control mice. We replicated this analytical design using our pipeline (i.e. 4 del vs 4 control group A; 4 dup vs 4 control group B). Additionally, we processed the data through our pipeline after combining all of the control samples (4 del vs 8 control A+B; 4 dup vs 8 control A+B). In both analytical designs we detect groups of genome-wide, non-CNV DEGs overlapping between deletion and duplication tissues (Figure S4). For the grouped-controls analysis, 51 non-CNV DEGs overlapped (Figure S4A). For the independent-controls analysis, we found 37 non-CNV DEGs that overlapped between the deletion mice and the duplication mice (Figure S4B).

PCSK9 expression levels were also evaluated in the mouse model RNA-Seq data. In both analyses (independent controls and grouped controls), we detected a similar trend of PCSK9 expression (Table S2). Namely, in the deletion mice PCSK9 expression was increased (.325 log2 fold change in independent-controls; .232 log2 fold change in grouped-controls) and decreased in the duplication mice (−.156 log2 fold change in independent controls; −.064 log2 fold change in grouped controls). The magnitude of change was much lower than in our iPSC-iN data and none of the gene expression fold changes in mouse reached statistical significance, but one should bear in mind that the RNA was extracted from whole mouse cortex tissue, i.e. consisting of multiple different cell types.

### PCSK9 knockdown leads to altered developmental phenotypes in zebrafish

To test whether mirror-image changes in PCSK9 expression alone may yield developmental phenotypes that could be relevant with regards to those seen in 16p11.2 patients, we employed the use of the zebrafish model. To mimic the levels of gene expression reduction and gene overexpression, respectively, observed in the 16p11.2 iPSC-iN model, a PCSK9 morpholino, together with a control morpholino, was used to achieve reduction in expression levels, and, both human and zebrafish PCSK9 mRNA were used to achieve gene overexpression. The morpholinos, and separately the mRNAs, were injected into zebrafish embryos at the single-cell stage and development was followed for 5 days post fertilization (dpf). We observed no detectable changes in the control and overexpression zebrafish while phenotypic alterations were observed in the morpholino knockdown zebrafish. The knockdown resulted in effects on embryonic and early brain development such that the zebrafish exhibited a marked curvature of the spine and a darkening of and reduction of brain tissue (Figure 4 A1). Additionally, these zebrafish had a reduced body length (3.85 ± 0.0016mm (Con MO) vs. 2.64 ± 0.0057mm (PCSK9MO) vs. 3.77 ± 0.0012mm (PCSK9 mRNA); p <0.05, Student’s t-test, *N*=3 independent tests) and an increased interocular distance (0.12 ± 0.00010mm (Con MO) vs. 0.40 ± 0.0036mm (PCSK9MO) vs. 0.12 ± 0.00010mm (PCSK9 mRNA); p <0.05, Student’s t-test, *N*=3 independent tests) resulting from neural degeneration and oedema when compared to the control and overexpression zebrafish (Figure S5).

**Figure 4:**
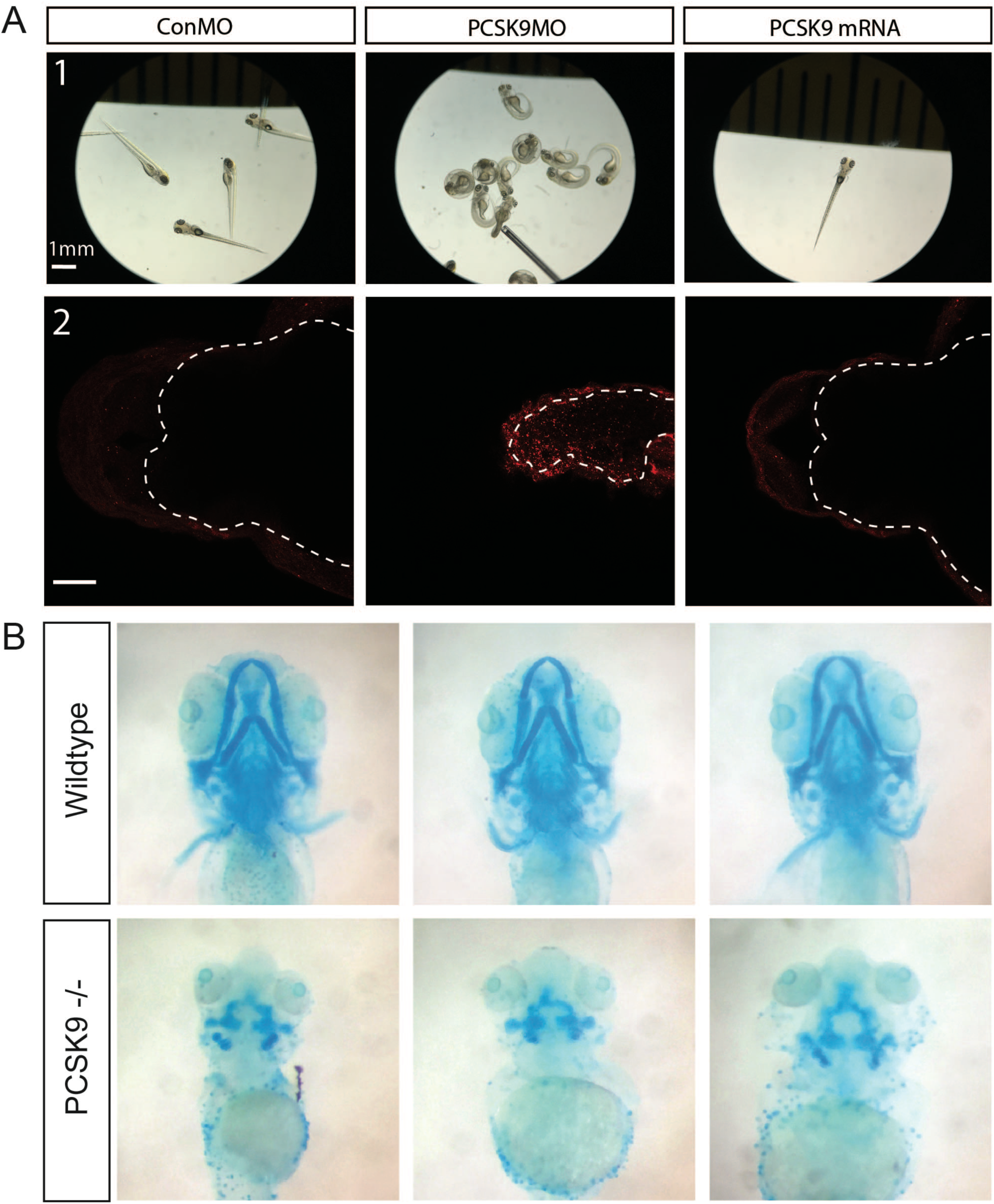
Reduction of PCSK9 expression in zebrafish impact development. **(A)** PCSK9 expression was altered in zebrafish by injecting a PCSK9 morpholino (PCSK9MO) to knockdown expression and PCSK9 mRNA to increase expression to comparable levels seen in the 16p11.2 CNV patients. A control morpholino (ConMO) was also injected to ensure no effects seen were from the morpholino injection. **(A1)** The knockdown resulted in defects on early brain/embryonic development; primarily the zebrafish had a curvature of the spine and a darkening and reduction of the brain tissue. By comparison, the control and overexpression zebrafish were unaffected. **(A2)** Caspase-3 immunostaining reveals that an increase in apoptosis is occurring in the 1dpf PCSK9 knockdown zebrafish within the hindbrain **(marked by dotted line)**. **(B)** At 5 dpf, Alcian Blue staining shows disruption of the ventral head structures including the jaw, presumptive operculum and pectoral fins of the PCSK9 −/− zebrafish which was not seen in the in-clutch wildtype zebrafish.

To determine the cause of the darkening and reduction of brain tissue, caspase-3 immunostaining was performed at 1 day post fertilization (dpf) and 5 dpf to test for an increase in apoptosis activity. At both days, we see an increase in apoptosis when compared to the other two zebrafish groups with more caspase-3 fluorescence at 1 dpf (Figure 4 A2).

To further test the effect of loss of PCSK9 on the development of zebrafish, we characterized the phenotype of heterozygous mutant crosses that contained PCSK9 +/+ PCSK9 +/− and PCSK9 −/− fish from 1dpf through 5dpf. While there were imperceptible differences between wildtype and heterozygous mutants, we detected at appropriate Mendelian ratios that PCSK9 −/− mutant fish (verified *post-hoc* by larval genotyping) exhibited similar developmental phenotypes in body plan and interocular distance as seen with morpholino knockdown, indicating the specificity of the aberrations with PCSK9 perturbation. Specifically, we observed reduced body size and curved body shape as well as small brains and eyes with increased interocular distance. Oedema was also observed in the heart, brain and around the eyes. The swim bladders of these fish were elongated. The effect of null mutation was embryonic lethal since fish with this genotype did not survive past 5-6dpf.

Prior to lethality, we detected a disruption of the development of ventral structures and further characterized these with Alcian Blue staining. Following staining of the cartilage at 5dpf, we observed abnormalities of the ventral head structures including the jaw, presumptive operculum and pectoral fins in the PCSK9 −/− zebrafish, which was not seen in the in-clutch wildtype zebrafish (Figure 4 B). Only mid-to-dorsal structures such as the otic vesicular cartilaginous structures remain in the PCSK9 −/− zebrafish. These results suggest that part of PCSK9’s function could be involved in the specification of ventral structures, failure of which, in complete absence, affect overall development with serious consequences to lifespan.

### DNA methylation altered genome-wide

Next we analyzed genome-wide patterns in DNA methylation marks in the CNV lines. A subset of the cell lines were (2 control lines, 2 deletion lines, 2 duplication lines) were analyzed for differentially methylated regions (DMRs) at both the iPSC and iN stages. We found similar numbers of DMRs ranging from 185 in iPSC-Dup to 286 in iN-Dup, with iPSC-Del (237 DMRs) and iN-Del (217 DMRs) in between (Figure 5 A). For each analysis, the DMRs were differentiated into hypermethylated or hypomethylated. In all four comparisons a majority of the DMRs were found to be hypermethylated. The lowest ratio of hypermethylated to hypomethylated is approximately 2:1 (iPSC-Del) and the highest ratio is 7:1 (iN-Dup). Within the CNV region, we see that cell stage does not impact the number of DMRs found, but CNV type has a large impact on DMR count genome-wide, with the duplication analyses having roughly twice the number of DMRs as the deletion analyses (iPSC-Dup 201, iN-Dup 236; iPSC-Del 98, iN-Del 96). A further breakdown of those numbers showed more of the DMRs being hypomethylated than hypermethylated, particularly in the duplication comparisons. iPSC-Del have a distribution of hyper/hypo of 41/57, iPSC-Dup show 89/112, iN-Del show 47/49, and iN-Dup show 83/153. However, very few of these DMRs passed a 0.1 minimum threshold for DNA methylation change with only 22 DMRs across all four comparisons passing this threshold (6 in iPSC-Del, 6 in iPSC-Dup, 2 in iN-Del, 8 in iPSC-Dup).

**Figure 5:**
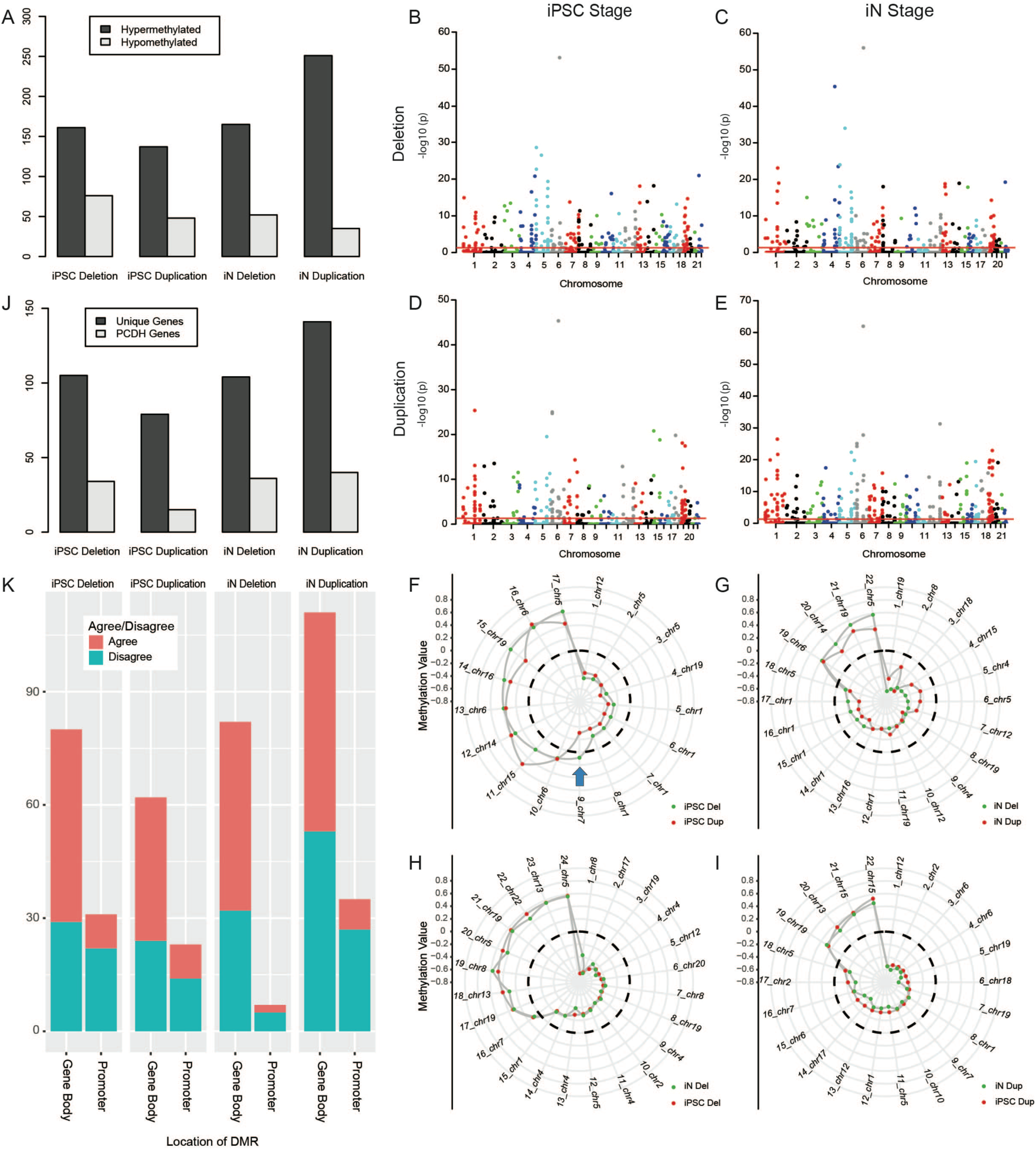
Global alterations to methylation from the presence of the CNV. The presence of the CNV has genome-wide effects on patterns of DNA methylation. **(A)** The total number of differentially methylated regions (DMRs) found in each analysis and the breakdown of DMRs found to be hypermethylated or hypomethylated compared to controls. **(B-E)** DMRs were found in each of the 4 analyses and the DNA methylation change associated –log10(p) was plotted by chromosomal location for each DMR, with the red line representing FDR 0.05 statistical cutoff. The values were plotted for the **(B)** iPSC deletion, **(C)** iN deletion, **(D)** iPSC duplication, and **(E)** iN duplication. **(F-I)** Overlapping DMRs from cell stage and CNV-type comparisons plotted by methylation value and labeled by a unique number value and chromosome location. Plots shown are DMRs overlapping **(F)** at the iPSC stage, **(G)** at the iN cell stage, **(H)** between cell stages for deletion patients, and **(I)** between cell stages for duplication patients. Dotted line represents zero change in methylation. **(J, K)** The DNA methylation and gene expression data were integrated to evaluate the potential effect of the epigenetic alterations on the gene expression changes. DMR/Gene overlaps were found based on the chromosomal location of the DMR and if the DMR was within the promoter region (+2000 bp of the gene’s start site) or gene body. **(J)** The number of unique genes with at least one DMR located within the gene’s promoter region or gene body. Also shown is the number of PCDH genes found within the unique genes lists. **(K)** When comparing the altered DNA methylation and gene expression changes, the comparisons were separated by location of the DMR and evaluated for DNA methylation + expression agreement. For the promoter region, the comparison was marked as an agreement if gene expression decreased while DNA methylation increased or gene expression increased while DNA methylation decreased. For the gene body, the opposite of the promoter agreement conditions meant the comparison was an agreement.

When examining the genomic locations of the DMRs (Figure 5 B-E) we found that, just as in the gene expression changes, the epigenetic alterations were spread across the genome and were not localized to chromosome 16 or enriched in the CNV locus. The number of DMRs found on each chromosome does not strictly track with chromosome size, number of protein coding genes per chromosome, or the number of CpG regions tested per chromosome. When grouped together by cell stage, chromosome 5 has the most DMRs (10.2% of all iPSC DMRs) followed by chromosomes 1 (10%) and 19 (9.5%). When grouped together by CNV type, a majority of the DMRs found on chromosome 5 (61 of 101) are contributed by the Del patient lines and a larger number of DMRs on chromosomes 1 and 19 (61 of 93, 53 of 89 respectively) are contributed by the Dup patient lines. Only 28 DMRs are located on chromosome 16, making it the chromosome with the 14^th^ most DMRs.

Similar to the gene expression analyses, we wanted to test which DMRs from the individual group analyses overlapped with one another. There were similar numbers of overlapping DMRs across the analyses: 17 overlapping for iPSC, 22 overlapping for iN, 24 overlapping for Del, 22 overlapping for Dup (Figure 5 F-I).

### Integration of gene expression and altered DNA methylation datasets

We correlated the genomic locations of the DMRs with those of all known genes to determine the extent to which they overlapped, which we refer to as DMR/gene overlaps (DGOs). When filtering for the DGOs that contain genes with detectable expression, we find 155 such DGOs in iPSC-Del, 99 in iPSC-Dup, 170 in iN-Del, and 173 in iN-Dup. Approximately 60% to 80% of the DGOs resided within a unique gene, with 106 of such unique genes in iPSC-Del, 78 in iPSC-Dup, 104 in iN-Del, and 138 in iN-Dup (Figure 5J). Interestingly, a large number of the genes are members of the PCDH family, where ∼19% to ∼35% of DGOs involve a PCDH gene.

Next, we analyzed whether the direction of DNA methylation change of a given DMR corresponded with the direction of gene expression change of the co-localized gene given, taking into account the genomic location of the DMR within the gene. If a DMR resides within the promoter region of a gene then hypermethylation of the DMR paired with reduction in gene expression is considered as agreeing (and vice versa). If a DMR resides within the gene body, the reverse constellation is considered as agreeing (Figure 5K). For a majority of DMRs located in promoter regions there was disagreement with the gene expression changes, with an almost 2:1 ratio of agreement to disagreement across all analyses. There was much more agreement between DNA methylation changes and gene expression level changes when DMRs are located within the gene body, with more than 50% of DGOs across all analyses showing agreement.

## Discussion

We were able to successfully grow and differentiate the iPSCs into iNs as verified by the immunostaining and WGCNA pathway analyses. This shows that the model used here is an efficient option for obtaining sufficient numbers of physiologically relevant cells in order to perform multiple functional genomics assays.

The large CNVs on chromosome 16p11.2 generally had the expected effect on expression levels for the genes that are encoded within the CNV boundaries, namely the expression changes follow the changes in gene copy number. In addition we observed that there is a marked genome-wide effect on gene expression levels, well beyond chromosome 16. There are two genes encoded within the CNV boundaries that act as transcription factors, *MAZ* and *TBX6*. This differential expression of two transcription factors could be the basis for at least some of the genome-wide expression level changes that we observed. However, an analysis of transcription factor binding motifs did not uncover an enrichment for the known binding motifs for *MAZ* and *TBX6* (data not shown), leaving this possible explanation for some of the global gene expression changes unconfirmed at this time. Many, if not all of the genes encoded within the CNVs can be expected to play potentially important roles in functional pathways.

Our reanalysis of the gene expression data from the 2014 Blumenthal et al. study using mouse models for both the deletion and duplication CNVs revealed a similar finding of overlapping non-CNV DEGs in the 16p11.2 mouse model. This supports the interpretation that this particular facet of transcriptional dysregulation associated with 16p11.2 copy number alterations is not exclusive to the iPSC model system. Reanalysis of the mouse data also strengthened support for the observations related to the expression changes seen for PCSK9.

Furthermore, regarding the existence of this ‘core’ set of overlapping DEGs genome-wide, it may be that these overlapping DEGs are at the core, on the molecular level, of the phenotypes that are seen in both deletion and duplication carriers, while the non-overlapping DEGs may be to a greater degree involved in the molecular basis of the phenotypes that are more commonly seen diverging in the patients.

The functional enrichment analyses identified enrichment in biological processes involving synaptic signaling, as well as neuronal components involving dendrites and somatodendritic compartments. This was in line with the findings of the 2017 Deshpande et al. study, reporting morphological changes in soma size, dendritic length, and synaptic density in the differentiated neurons of 16p11.2 CNV patients. While the gene expression changes were not opposing, unlike the reported morphological changes, this still provides further indirect biological validation for our genomic findings.

Our observation of opposite-direction gene expression level changes for PCSK9 implies a potential role of this gene in the etiology of the opposing phenotypes of the deletion and duplication patients. PCSK9 could be contributing to both the head-circumference (Norata et al., 2016) and body-mass-index phenotypes (Schulz & Schluter 2017), also considering the reduction in brain mass and body length seen in some of the zebrafish experiments. Such reductions were also consistent to those seen before with PCSK9 knockdown in zebrafish (Poirier et al., 2006). Our data also indicate that changes in PCSK9 expression levels at critical stages of development may also be playing a role in the development of facial abnormalities seen in 16p11.2 patients, for example the observed jaw malformations. As noted by Shinawi et.al., “Subjects with the deletions shared the following features: broad forehead, micrognathia, hypertelorism, and a flat midface. The broad forehead, macrocephaly and flat midface give these patients a distinct facial gestalt.” In our zebrafish experiments, we observed that a reduction of PCSK9 expression levels resulted in major disorganization and underdevelopment of ventral jaw structures. One needs to bear in mind, however, that PCSK9 in humans plays a smaller role in head and facial development than in zebrafish, and much further work in this context is needed, also in the light of reported that PCSK9 knockout mouse models do not exhibit facial or head circumference abnormalities (Rashid et al., 2005; Seidah et al., 2008).

The CNVs on 16p11.2 alter DNA methylation patterns similarly in both deletion and duplication patients as well as at both differentiation stages. Overall, the CNVs increase levels of DNA methylation across the genome except within the CNV region itself, where there is little to no significant change in DNA methylation levels. Chromosome 16 as a whole is also not disproportionately impacted on the level of DNA methylation. As there is generally a similar genome-wide impact on DNA methylation in all four analyses, it could be that the deletion and duplication of the 16p11.2 region are affecting the same or similar DNA methylation regulation pathways.

We did not find a strong correlation between the altered DNA methylation patterns and specific changes in gene expression in this study. However, the protocadherin (PCDH) gene family stood out as a candidate for follow-up study. Dysregulation of the protocadherins on both the levels of epigenetic marks and gene expression has been found to be associated with human disease, including neurological disorders. The review by Hajj et. al 2017 outlines the known associations that these dysregulations have with disorders such as schizophrenia (Narayanan et al., 2015; Gregrio et al., 2009), bipolar disorder (Gregrio et al., 2009), and ASD (Bucan et al., 2009; Morrow et al., 2008). We did not observe altered expression for the PCDH genes. At the same time, PCDH expression levels overall were very low in the early neuronal cell type that is being replicated by the iN cells in this study.

With this study we have demonstrated that by using a disease relevant cell type, and the precursor cells to this relevant cell type, we are able to better identify genes and pathways that may play an etiological role in the development of the disease phenotype. Studying the cells as they progress through the differentiation process may allow us to attain a more complete picture of the genetic and epigenetic changes occurring as a result of the presence of the large CNVs on 16p11.2. For example, PCSK9 has low levels of expression in a mature neuronal cell type but nevertheless an early perturbation of PCSK9 expression levels at the stem cell stages could have a profound effect on nervous system development, for example if it leads to abnormally high or low levels of apoptosis at critical periods of development.

The availability of the iPSC lines used in this study from a public cell repository means that these lines, as well as the functional genomics data generated by us for these lines, constitute an important resource for the further study and eventual understanding of the molecular basis of neuropsychiatric, neurodevelopmental phenotypes such as those observed in many individuals with CNVs in 16p11.2.

## Author Contributions

Thomas R. Ward - conceptualization, methodology, validations, formal analysis, investigation, writing, visualization.

Xianglong Zhang - methodology, validations, formal analysis, investigation, visualization

Louis C. Leung - validations, formal analysis, investigation, visualization

Bo Zhou- writing – review & editing

Kristin Muench - methodology, investigation

Arineh Khechaduri - investigation

Melanie J. Plastini - investigation, validations

Carol Charlton - methodology

Reenal Pattni – methodology, validations

Steve Ho - validations

Marcus Ho - methodology

Yiling Huang - methodology

Joachim F. Hallmayer – supervision, funding acquisition

Phillippe Mourrain – conceptualization, supervision

Theo D. Palmer – conceptualization, supervision, funding acquisition

Alexander E. Urban- conceptualization, supervision, results interpretation, writing, funding acquisition

## Acknowledgements

This work was supported by grants to TDP (NIH/NIMH R01MH108659) and AEU (NIH DP2 MH100010-01). AEU was also supported by funds from the Tashia and John Morgridge Faculty Scholar program and by funds from Bruce Blackie.

## Supplementary Figures

**Figure S1, Related to Figure 1:**
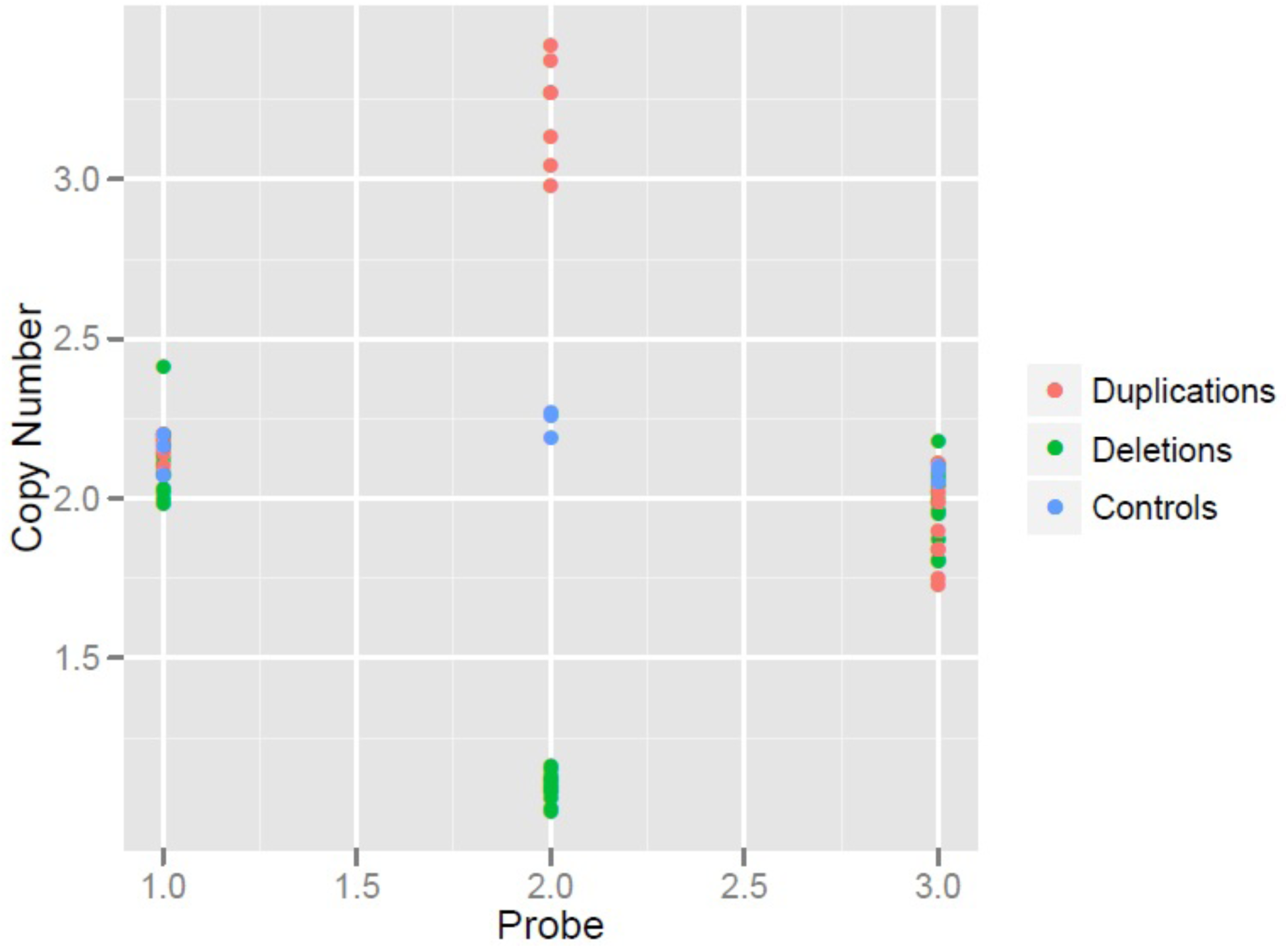
ddPCR verification of the deletion and duplication of the 16p11.2 region. Using TaqMan Copy Number Assays, a ddPCR experiment was run on the 12 patient samples at the iPSC stage to verify the presence of the CNV in the patient samples. The control samples were included as a positive control for a copy number of two for the region. Three assays were used: one located within the boundaries of the CNV region (Probe #2) and two located outside of the boundaries to serve as controls (Probes #1 and 3). The respective copy numbers of each probe are plotted for each sample.

**Figure S2, Related to Figure 1:**
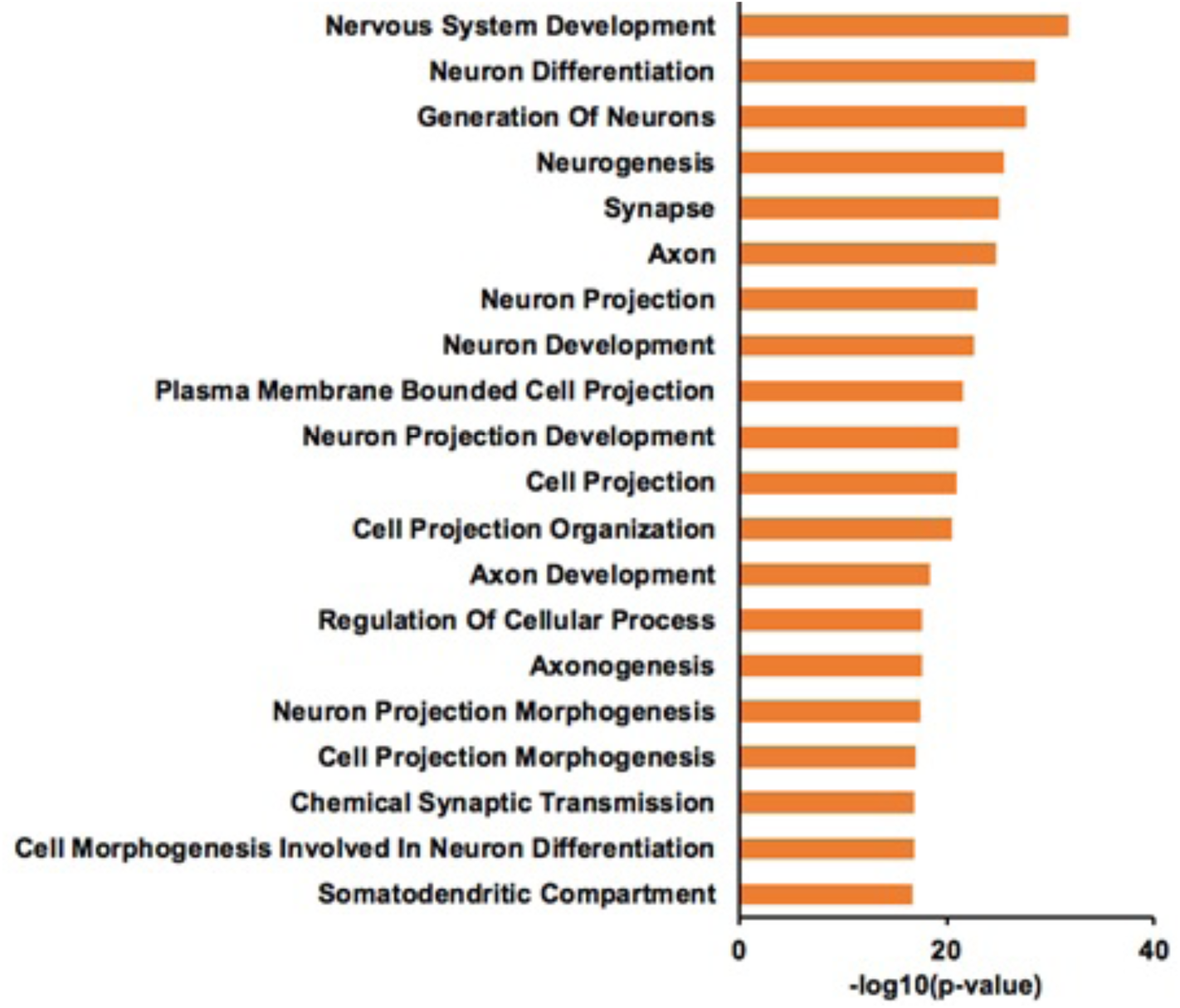
Pathway analysis of upregulated genes at the iN stage. WGCNA pathway analysis was performed by comparing the iPSC stage to the iN stage for all cell samples. The analysis identified 6,604 genes that have increased expression in the iN cells compared to the iPSCs. The upregulated gene pathways are plotted with their corresponding enrichment p-value, converted to –log10(p-value) for the plot.

**Figure S3, Related to Figure 3:**
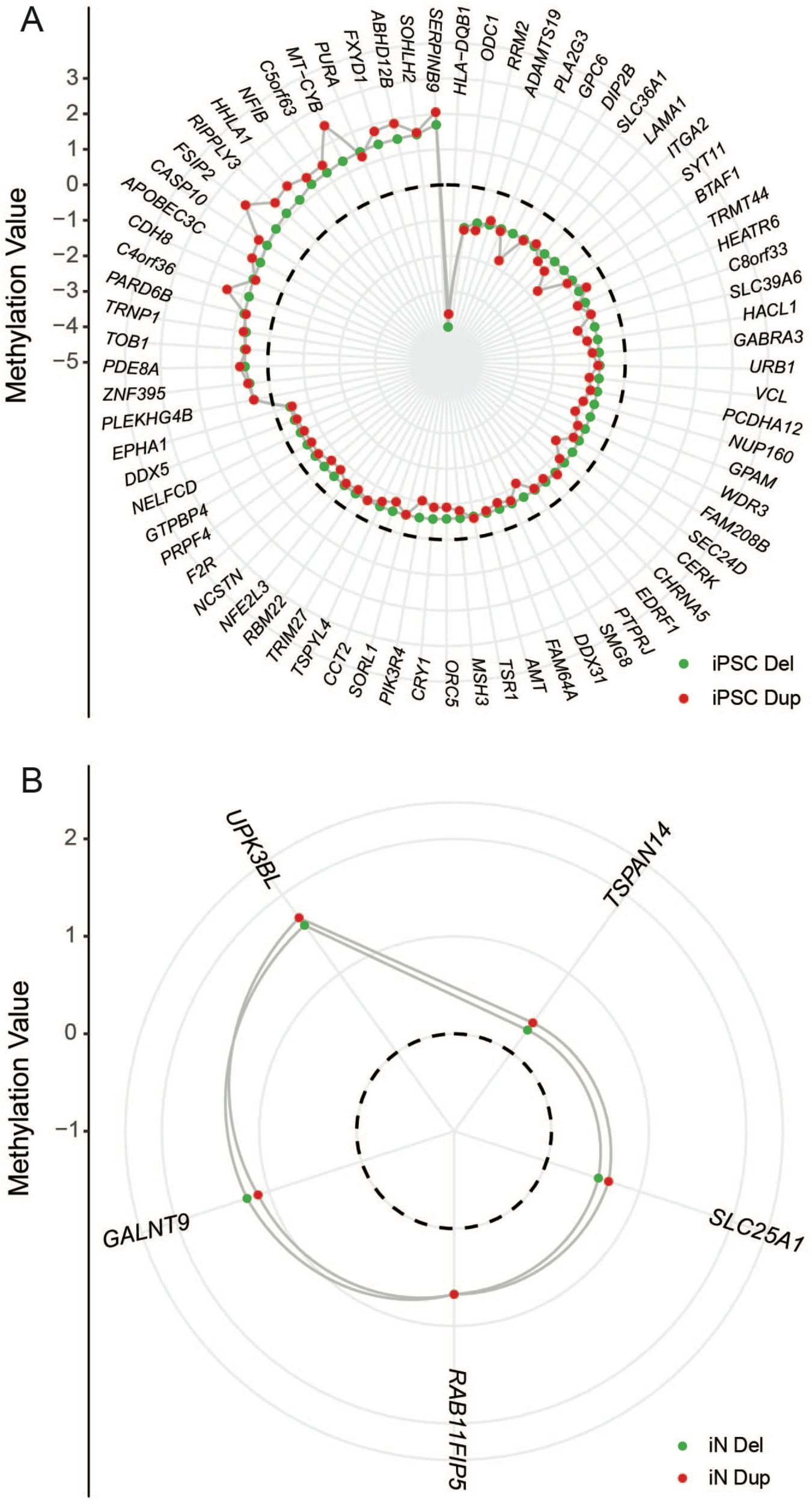
Overlapping DEGs between cell stage analyses using DESeq2 data. The four analyses done in Cuffdiff were repeated using an alternative pipeline, DESeq2. With this data, all DEGs that had a FDR ≤ .05 were compared between deletion and duplication patients at both cell stages resulting in **(A)** 73 overlapping DEGs at the iPSC stage and **(B)** 5 DEGs overlapping at the iN stage. The log2 fold change values for each DEG were plotted from each analysis, green representing the deletion analysis values and the red representing the duplication analysis values. Dotted line represents zero fold change difference in expression.

**Figure S4, Related to Figure 3:**
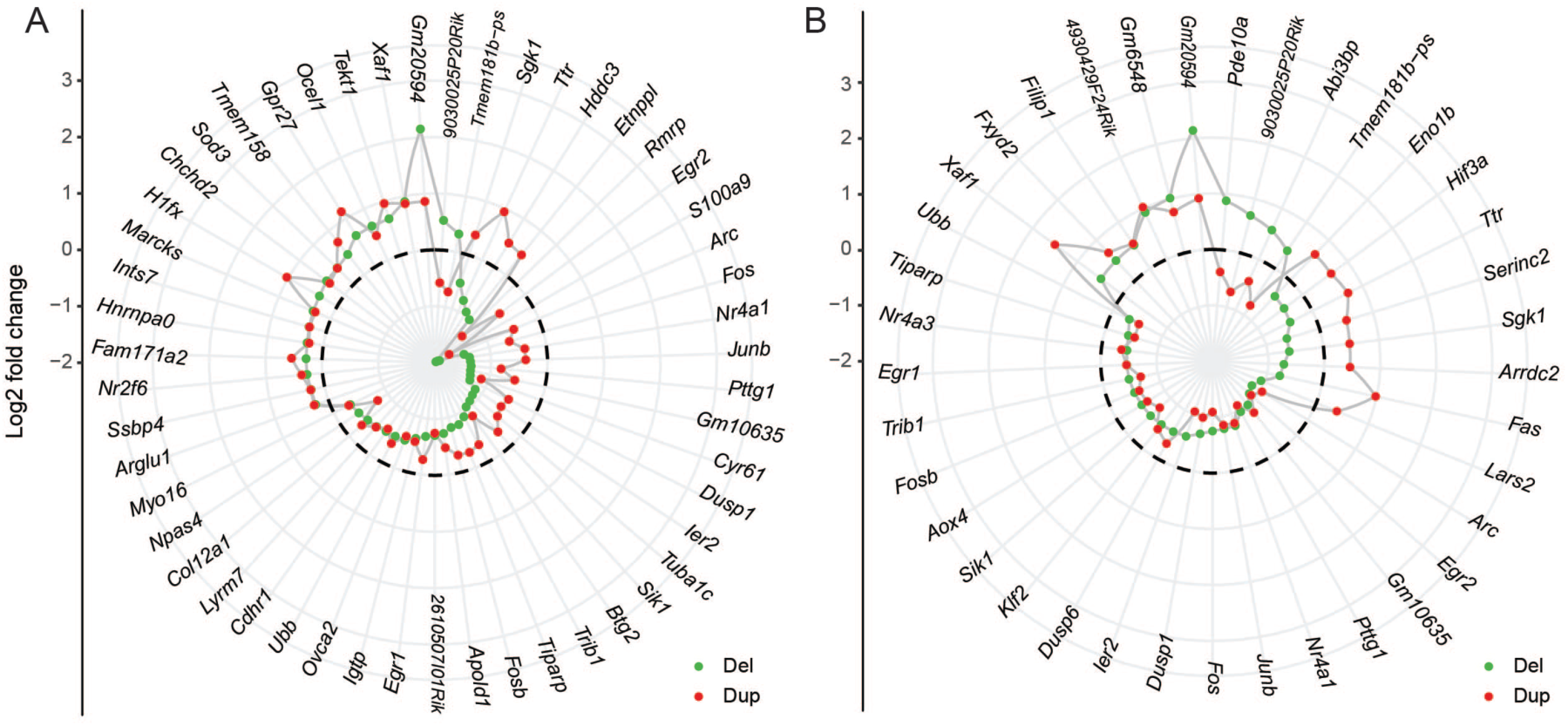
Mouse brain RNA-Seq data radar plots. The 16p11.2 CNV mouse cortex RNA-seq data acquired from 2014 Blumenthal et al. was analyzed using the same analysis pipeline as the 16p11.2 CNV patient samples. We replicated the experimental design with independent controls (4 del vs 4 control group A; 4 dup vs 4 control group B) as well as combining all control samples (4 del vs 8 control A+B; 4 dup vs 8 control A+B). Overlapping DEGs were compared for the **(A)** grouped controls analyses and the **(B)** independent controls analyses. The log2 fold change values were plotted for each DEG, green representing the deletion analysis values and the red representing the duplication analysis values. Dotted line represents zero fold change difference in expression. **(A)** 45 of the 51 overlapping DEGs in the grouped analysis and **(B)** 25 of the 37 overlapping DEGs in the independent analysis had fold changes in the same direction.

**Figure S5, Related to Figure 4:**
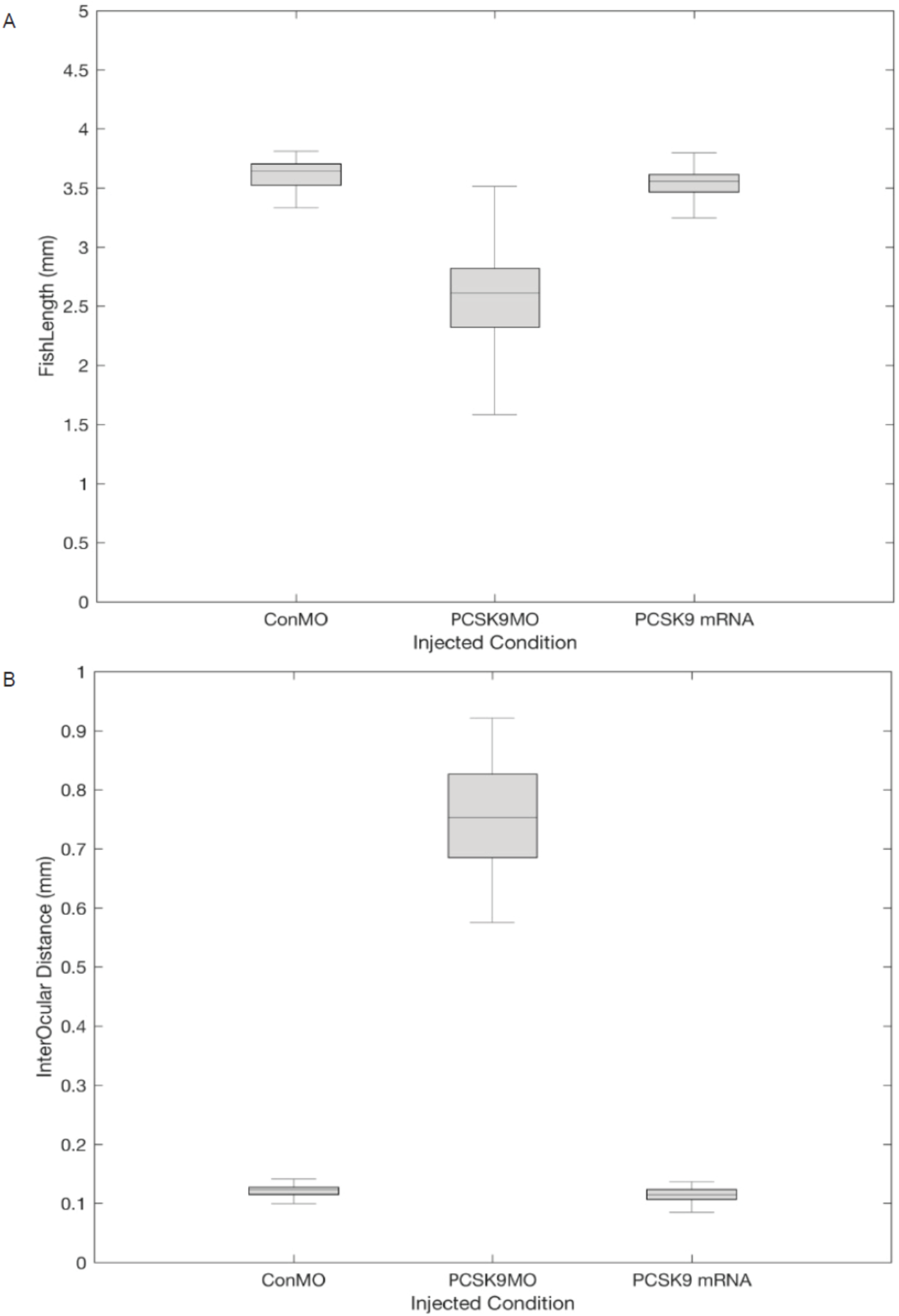
Altered length and interocular distance in the PCSK9MO zebrafish. Boxplots of the **(A)** fish length and **(B)** interocular distance of each of the zebrafish groups.

## Methods Details

### Cell culture and differentiation

#### Cell lines

A total of 14 16p11.2 CNV patient iPSC lines (9 deletion and 5 duplication) were obtained from the Simons Variation in Individuals Project (Simons VIP; https://www.sfari.org/resource/simons-vip/). Each patient currently has multiple clonal lines available. Twelve clonal cell lines from 6 patients carrying the 16p11.2 CNV (3 deletion, 3 duplication, and 2 independently derived clones from each patient) were used. These lines were paired with 4 independent control lines that were matched for age, sex, and ethnicity. Cell line IDs as used by Simons VIP:

Deletions: 14758×3 101.7 & 101.8, 14765×2 101.1 & 101.2, 14781×16 101.3 & 101.4

Duplications: 14723×10 202.7 & 202.8, 14756×9 201.1 & 201.2, 14756×16 101.1 & 101.2

Controls: 5401.1, 6632.4, 726.1, 8738.3

#### Cell culture

Each iPSC line was cultured using mTeSR media (Stemcell Technologies, 85850) in clear plastic BioLite 6-well multidish plates (Thermo Scientific, 130184) coated with Matrigel hESC-qualified Matrix (BD Biosciences, 356234). The cells were kept at 37° C and 5% CO_2_; media was changed daily. The cells were passaged every 4 to 5 days once the wells reached ∼ 75-100% confluency. For passaging, cells were washed with DPBS (Life Technologies, 14190-144), dissociated with TrypLE Express Enzyme (1X) (ThermoFisher, 12604013) and added to fresh media on a new plate and/or added to freezing media for storage. All lines were grown concurrently to avoid batch effects. One duplication line (14723×10 202.7) was not viable due to large amounts of cell differentiation and was not included in the analysis. New starter cultures of this line have been made and are shown to be viable.

#### Neuronal differentiation

All iPSC cell lines were differentiated to induced neurons (iNs) following the two-week protocol described by Zhang et al. Once the cells reached ∼80-100% confluency, the wells were split at a 1:3 ratio using Accutase (Stem Cell Technologies, 7920) for single cell density plating. One day post-split, a single well was transfected with 2 uL of each of the 3 viral vectors acquired from the Genome Virus and Vector Core (GVVC) at Stanford University. The transfected cells were then subjected to puromycin selection and monitored for 7 days. On day 7, the immature iNs were collected and underwent DNA/RNA extraction. The protocol was modified to be stopped at day 7 instead of the complete two-week protocol to obtain immature cortical glutamatergic neurons. GFP fluorescence was used to verify successful integration of the vectors and MAP/TUB staining was done to ensure complete differentiation into neuronal cells (Supp Fig 1).

#### DNA/RNA extraction

DNA and RNA were extracted using the All Prep kit (Qiagen, Cat. #80204) once the wells reached ∼80-100% confluency (∼1 to 3 million cells). mRNA purification was performed using the Dynabeads mRNA Purification Kit (Invitrogen, Cat #61006) for RNA sequencing.

### Sequencing analyses

#### Short-Insert Whole-Genome Library Preparation

iPSC DNA was sheared to 400bp on a Covaris E210R (Covaris, Woburn, MA). Libraries were constructed with 500ng of input DNA and created according to standard ligation methods using the Kapa Hyper Prep Kit (Kapa Biosystems, Wilmington, MA). Each library was amplified with 3 PCR cycles and size selected between 300 and 700 bp using field-inversion gel electrophoresis on the Pippin Pulse System (Sage Science, Beverly, MA) and fragment sizes were verified on a Bioanalyzer 2100 (Agilent Technologies, Santa Clara, CA). Libraries were sequenced on an Illumina Next-seq 500 using a NextSeq 500/550 High Output v2 kit (300 cycles) (Illumina, FC-404-2004).

#### RNA sequencing and analysis

The libraries for all samples at both iPSC and iN stages were prepared using the NEBNext Ultra RNA Library Prep Kit for Illumina (New England BioLabs, E7530S) and were sequenced on the Illumina NextSeq 500 using a NextSeq 500/550 High Output v2 kit (Illumina, FC-404-2002) (2X75 bp) for the iPSCs and a NextSeq 500/550 High Output v2 kit (Illumina, FC-404-2004) (2X150 bp) for the iNs. The 2×150 bp reads were trimmed to 2×75 bp for more direct comparison with the iPSC reads in the analysis. Reads were first mapped to hg38 (Y chromosome excluded) (GCF_000001405.36) using TopHat2 (v2.0.9) (Kim D et al., 2013) with transcript index built from GENCODE comprehensive gene annotation (release 23) using ‘‘*--library-type fr-firststrand*’’ and ‘‘*-- transcriptome-index*’’.

Differential expression was performed using Cuffdiff (v2.1.1) (Trapnell et al.) with “-- dispersion-method per-condition --library-type fr-firststrand’’ against the GENCODE comprehensive gene annotation. Only the genes with FPKM > .05 were used for downstream analysis. Deletion and duplication samples were individually compared with control samples at both iPSC and iN stages. Four separate analyses were performed: control samples versus deletion samples at iPSC stage (iPSC Del), control samples versus duplication samples at iPSC stage (iPSC Dup), control samples versus deletion samples at iN stage (iN Del), and control samples versus duplication samples at iN stage (iN Dup).

Given the heterogenous genetic background of the CNV patients, to see if our differential expression results were consistent in a distinct testing framework we elected to use a secondary analysis pipeline. Cutadapt ^1^ (version 1.8.1) was used to trim Illumina TruSeq adapters and low-quality ends from the raw reads. Bowtie2 ^2^ (version 2.3.1) was used to align the trimmed reads to the GENCODE comprehensive gene annotation (release 23, hg38) and RSEM ^3^ (version 1.2.30) was used to quantify gene expression in a strand-specific manner by setting parameter “--forward-prob 0”. DESeq2 ^4^ (version 1.12.4) was used to perform differential expression analysis. Genes with FDR adjusted *p*-value < 0.1 were considered to be significant for iNs and < 0.05 for iPSCs. Only the protein-coding genes with detectable expression (FPKM > 1) were used for differential expression analysis, WGCNA and pathway enrichment analysis.

To identify modules of co-expressed genes, we performed weighted-gene coexpression network analysis (WGCNA, version 1.46) ^5^ using all the protein-coding genes with detectable expression (approximately 16,000 genes). Normalized count data obtained by the “varianceStabilizingTransformation” function in DESeq2 were used. Signed network was constructed for each comparison.

#### qPCR verification for gene expression

qPCR primers for PCSK9, GAPDH, and ALDOA was found using RTPrimerDB (www.rtprimerdb.org) and PrimerBank (https://pga.mgh.harvard.edu/primerbank/). cDNA was synthesized from the mRNA of the samples using ReadyScript cDNA Synthesis Mix (Sigma-Aldrich) and the qPCR was done using KAPA SYBR FAST qPCR Master Mix and the corresponding protocol. The qPCR was run on a QuantStudio 6 Flex Real-Time PCR System (ThermoFisher Scientific).

#### ddPCR verification of the 16p11.2 CNV

TaqMan Copy Number Assays (ThermoFisher Scientific) were used in a ddPCR experiment prepared and run on a Bio Rad QX 200 Droplet Reader (Bio Rad). Three different TaqMan assays were used, one was located within the CNV region (location chr16:29901542, Cat. ID Hs02040751_cn) and two other assays (location chr16:29228600, Cat. ID Hs05451406_cn; location chr16:30389709, Cat. ID Hs05421015_cn) each located on either side of the CNV boundaries.

#### CpG-Capture bisulfite sequencing and analysis

Bisulfite converted libraries were prepared using the SeqCap Epi CpGiant Probes kit (Roche, Catalog No. 07138881001) following the manufacturer’s protocol. For this analysis, six samples at the iPSC and iN cell stages were included: two control samples (6632.4 & 726.1), two deletion samples (14723×10 202.8 & 14756×9 201.2), and two duplication samples (14758×3 101.7 & 14765×2 101.2). These samples were sequenced on the Illumina Nextseq 500 using the NextSeq 500/550 High Output v2 kit (300 cycles) with 30% PhiX, generating an average of 90 million reads generated for each sample. After trimming the adapters and low-quality ends using Cutadapt (Version 1.8.1, Marcel Martin), the reads were mapped to GRCh38.p10 (GCF_000001405.36) using Bismark (Version 0.16.3; Krueger and Andrews). Duplicates were removed by the deduplicate_bismark script in the Bismark package. In cases where the 3’-end sequences of the paired-end reads overlap due to the size of the library insert, only one copy of the overlapping region was retained, after clipping the read with the lower average read quality in the overlap region using the “*clipOverlap*” tool in bamUtil (Version 1.0.14; https://github.com/statgen/bamUtil). On-target read rate and coverage were calculated by Qualimap (Version 2.1; Garcia-Alcalde et al).

Methylation ratio for each CpG was extracted using the bismark_methylation_extractor script in the Bismark package. For each sample, only CpGs covered with at least ten reads were included in the downstream analysis. Differentially methylated regions (DMRs) were identified between the iPSC and iN cell types as well as between patients and controls within each cell type using *metilene* (Version 0.2-6; Jühling et al.) with ≥ 3 CpGs and a mean methylation difference between the two compared groups of ≥ 0.2. DMRs, and a FDR corrected *p*-value < 0.05 was considered significant.

### Zebrafish experiments

#### PCSK9 Plasmids, morpholinos and mutants

Full length PCSK9 cDNAs for zebrafish (synthesized by Genscript) and human (Dharmacon) were separately cloned into the pCS2+ backbone (Addgene #17095) with EcoRI and XbaI restriction enzyme sites. Plasmids were linearized with NotI restriction enzyme (NEB) and *in vitro* transcription was performed with mMessage machine SP6 Kit (Ambion) according to manufacturer’s instructions. Capped mRNA were purified with the RNAeasy kit (Qiagen). Standard scrambled control (5’-CCTCTTACCTCAGTT-ACAATTTATA-3’), mismatched (5’-GAGGCTTGTCATTGTCTCTGGTTTC-3’) and antisense (5’-GACGCTTCTCATTCTCTGTGCTTTC-3’) morpholinos were designed as previously described (Poirier et al., 2006). The PCSK9 heterozygous mutant with a C > A premature stop codon in Tyr585 (sa23639, Zebrafish Mutation Project) was genotyped with the following primers (PCSK9 For: CATCTGGAGAAGCATCAGACTCTG; PCSK9 Rev: CCATTTACGTGACGTCACACTTAAC). In-crosses of heterozygous parents were used due to homozygous embryonic lethality.

#### Micro-injection

To alter the expression of PCSK9, 1-cell-stage wildtype zebrafish (leopard background) were injected through pulled glass needles by pulsatile injection using a Picospritzer (FemtoJet, Eppendorf). 50-100ng of full length zebrafish or human PCSK9 capped mRNA were injected for PCSK9 over-expression and 250μM of control, missense and antisense morpholino oligos were injected for PSCK9 knockdown. Embryos from the same clutch were used to prevent inter-clutch variations.

#### Caspase 3 staining

Wholemount Caspase-3 immunostainings were performed on 1 and 5 days-post-fertilization (dpf) zebrafish as previously described (Weber et al., 2016, Development) using an anti-caspase3 antibody (ab13847, Abcam). Embryos at 1 dpf were fixed in 4%PFA/PBS 6 h after treatment with EtOH. Larvae at 5 dpf were fixed after 4OHT treatment in 4%PFA/PBS for 3 h or overnight.

#### Image acquisition and analysis

Three independent experimental replicates were performed. Larvae were fixed in 4%PFA/PBS at 5dpf to perform body length and interocular distance analysis using these previously established developmental hallmark statistics (Escamilla et al., 2017, Nature). Larvae were photographed with a 12MP camera (iPhone6, Apple) with a 10xlens adaptor (Amazon.com) on a dissecting microscope (Zeiss) with a micrometer marker. Body length (from snout to tail fin) and interocular distance (distance measured between the eyes looking at the top-view of the zebrafish) were quantified with images imported into Fiji (ImageJ, NIH), distance metrics were measured and normalized to the in-image micrometer distance. Measurements were exported to Excel (Microsoft) and statistics were derived with custom MatLab (Mathworks) scripts. Analysis scripts are available upon request.

#### Cartilage staining

To study the disruption of the zebrafish ventral jaw structures, the cartilage was stained by Alcian Blue where 5dpf larvae were fixed in 4% paraformaldehyde overnight at 4 degrees C, followed by two rinses in 1x PBS Tween (0.1%) before staining overnight in Alcian Blue solution in EtOH (Electron Microscopy Sciences #26116-06) before washing and mounting in glycerol. Photographs were acquired as described above and were cropped using Photoshop (Adobe); otherwise presented unchanged.

## Supplementary Tables

**Table S1, Related to Figure 3:**
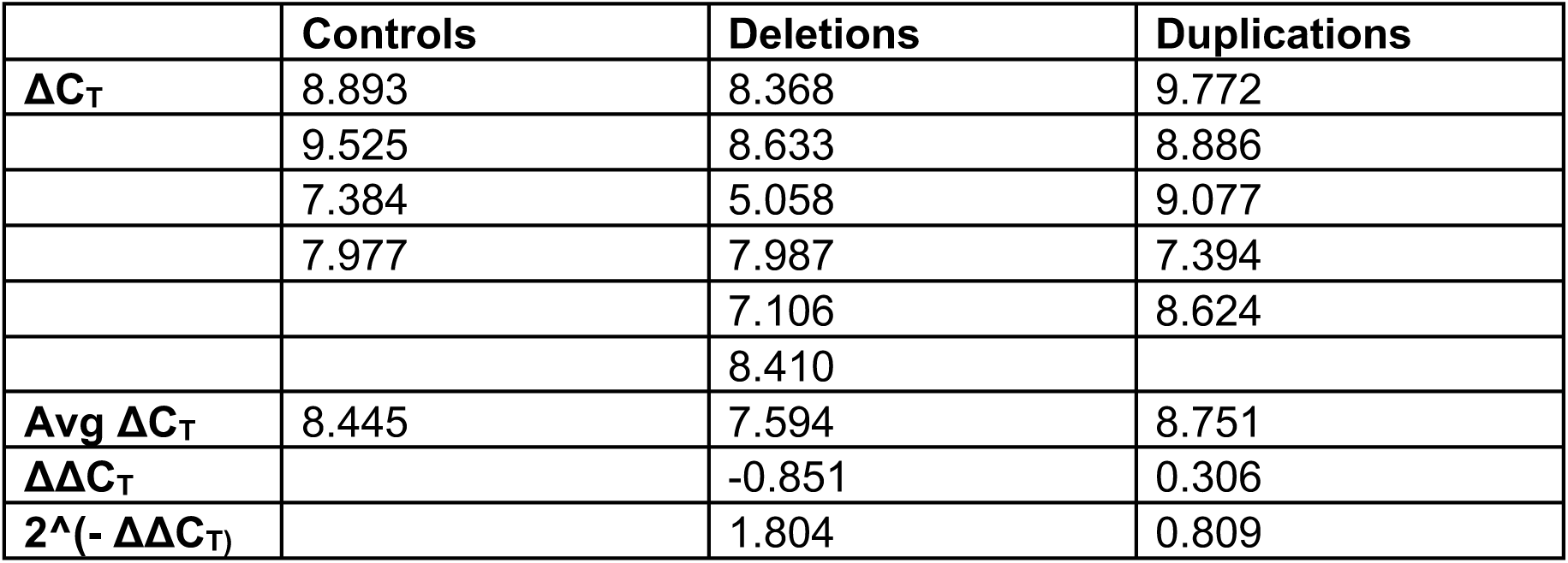
qPCR calculations for PCSK9 expression. Calculations for the qPCR experiment to find the expression changes of PCSK9 in the patient cells at the iPSC stage. A qPCR experiment was done using GAPDH as the control gene and ALDOA as a comparative gene with a known change in expression. ΔC_T_ calculations were made between the GAPDH values and PCSK9 values.

**Table S2, Related to Figure 3:** Mouse brain RNA-Seq data PCSK9 values. The FPKM, fold change, and p-values for PCSK9 in the CNV mouse data.

|  | **Control FPKM** | **CNV FPKM** | **Fold Change** | **P-value** |
| --- | --- | --- | --- | --- |
| **Del**<br>**(Independent)** | .662719 | .830143 | .324963 | .11315 |
| **Dup**<br>**(Independent)** | .758187 | .680505 | -.155946 | .4468 |
| **Del (Group)** | .709035 | .8326 | .231767 | .2026 |
| **Dup (Group)** | .709035 | .678298 | -.0639365 | .7311 |

## Literature References

Anders S, Pyl PT, Huber W. (2015). HTSeq--a Python framework to work with high-throughput sequencing data. Bioinformatics 31, 166–9.

Arbogast T, Ouagazzal AM, Chevalier C, Kopanitsa M, Afinowi N, Migliavacca E, Cowling BS, Birling MC, Champy MF, Reymond A, Herault Y. (2016). Reciprocal Effects on Neurocognitive and Metabolic Phenotypes in Mouse Models of 16p11.2 Deletion and Duplication Syndromes. PLoS Genet 12, e1005709.

Bernier R, Hudac CM, Chen Q, Zeng C, Wallace AS, Gerdts J, Earl R, Peterson J, Wolken A, Peters A, Hanson E, Goin-Kochel RP, Kanne S, Snyder LG, Chung WK; Simons VIP consortium. (2017). Developmental trajectories for young children with 16p11.2 copy number variation. Am J Med Genet B Neuropsychiatr Genet 174, 367–380.

Bijlsma EK, Gijsbers AC, Schuurs-Hoeijmakers JH, van Haeringen A, Fransen van de Putte DE, Anderlid BM, Lundin J, Lapunzina P, Pérez Jurado LA, Delle Chiaie B, Loeys B, Menten B, Oostra A, Verhelst H, Amor DJ, Bruno DL, van Essen AJ, Hordijk R, Sikkema-Raddatz B, Verbruggen KT, Jongmans MC, Pfundt R, Reeser HM, Breuning MH, Ruivenkamp CA. (2009). Extending the phenotype of recurrent rearrangements of 16p11.2: deletions in mentally retarded patients without autism and in normal individuals. Eur J Med Genet 52, 77–87.

Blumenthal I, Ragavendran A, Erdin S, Klei L, Sugathan A, Guide JR, Manavalan P, Zhou JQ, Wheeler VC, Levin JZ, Ernst C, Roeder K, Devlin B, Gusella JF, Talkowski ME. (2014). Transcriptional consequences of 16p11.2 deletion and duplication in mouse cortex and multiplex autism families. Am J Hum Genet 94, 870–83.

Bucan M, Abrahams BS, Wang K, Glessner JT, Herman EI, Sonnenblick LI, Alvarez Retuerto AI, Imielinski M, Hadley D, Bradfield JP, Kim C, Gidaya NB, Lindquist I, Hutman T, Sigman M, Kustanovich V, Lajonchere CM, Singleton A, Kim J, Wassink TH, McMahon WM, Owley T, Sweeney JA, Coon H, Nurnberger JI, Li M, Cantor RM, Minshew NJ, Sutcliffe JS, Cook EH, Dawson G, Buxbaum JD, Grant SF, Schellenberg GD, Geschwind DH, Hakonarson H. (2009). Genome-wide analyses of exonic copy number variants in a family-based study point to novel autism susceptibility genes. PLoS Genet 5, e1000536.

Buenrostro JD, Wu B, Chang HY, Greenleaf WJ. (2015). ATAC-seq: A Method for Assaying Chromatin Accessibility Genome-Wide. Curr Protoc Mol Biol 109, 21.29.1–9.

Chen J, Bardes EE, Aronow BJ, Jegga AG. (2009). ToppGene Suite for gene list enrichment analysis and candidate gene prioritization. Nucleic Acids Research doi: 10.1093/nar/gkp427.

Deshpande A, Weiss LA. (2018). Recurrent reciprocal copy number variants: Roles and rules in neurodevelopmental disorders. Dev Neurobiol 78, 519–530.

El Hajj N, Dittrich M, Haaf T. (2017). Epigenetic dysregulation of protocadherins in human disease. Semin Cell Dev Biol 69, 172–182.

Escamilla CO, Filonova I, Walker AK, Xuan ZX, Holehonnur R, Espinosa F, Liu S, Thyme SB, López-García IA, Mendoza DB, Usui N, Ellegood J, Eisch AJ, Konopka G, Lerch JP, Schier AF, Speed HE, Powell CM. (2017). Kctd13 deletion reduces synaptic transmission via increased RhoA. Nature 551, 227–231.

Fernandez BA, Roberts W, Chung B, Weksberg R, Meyn S, Szatmari P, Joseph-George AM, Mackay S, Whitten K, Noble B, Vardy C, Crosbie V, Luscombe S, Tucker E, Turner L, Marshall CR, Scherer SW. (2010). Phenotypic spectrum associated with de novo and inherited deletions and duplications at 16p11.2 in individuals ascertained for diagnosis of autism spectrum disorder. J Med Genet 47, 195–203.

García-Alcalde F, Okonechnikov K, Carbonell J, Cruz LM, Götz S, Tarazona S, Dopazo J, Meyer TF, Conesa A. (2012). Qualimap: evaluating next-generation sequencing alignment data. Bioinformatics 28, 2678–9.

Ghebranious N, Giampietro PF, Wesbrook FP, Rezkalla SH. (2007). A novel microdeletion at 16p11.2 harbors candidate genes for aortic valve development, seizure disorder, and mild mental retardation. Am J Med Genet A 143A, 1462–1471.

Girirajan S, Rosenfeld JA, Coe BP, Parikh S, Friedman N, Goldstein A, Filipink RA, McConnell JS, Angle B, Meschino WS, Nezarati MM, Asamoah A, Jackson KE, Gowans GC, Martin JA, Carmany EP, Stockton DW, Schnur RE, Penney LS, Martin DM, Raskin S, Leppig K, Thiese H, Smith R, Aberg E, Niyazov DM, Escobar LF, El-Khechen D, Johnson KD, Lebel RR, Siefkas K, Ball S, Shur N, McGuire M, Brasington CK, Spence JE, Martin LS, Clericuzio C, Ballif BC, Shaffer LG, Eichler EE. (2012). Phenotypic heterogeneity of genomic disorders and rare copy-number variants. N Engl J Med 367, 1321–31.

Girirajan S, Dennis MY, Baker C, Malig M, Coe BP, Campbell CD, Mark K, Vu TH, Alkan C, Cheng Z, Biesecker LG, Bernier R, Eichler EE. (2013). Refinement and discovery of new hotspots of copy-number variation associated with autism spectrum disorder. Am J Hum Genet 92, 221–37.

Green Snyder L, D’Angelo D, Chen Q, Bernier R, Goin-Kochel RP, Wallace AS, Gerdts J, Kanne S, Berry L, Blaskey L, Kuschner E, Roberts T, Sherr E, Martin CL, Ledbetter DH, Spiro JE, Chung WK, Hanson E; Simons VIP consortium. (2016). Autism Spectrum Disorder, Developmental and Psychiatric Features in 16p11.2 Duplication. J Autism Dev Disord 46, 2734–48.

Gregório SP, Sallet PC, Do KA, Lin E, Gattaz WF, Dias-Neto E. (2009). Polymorphisms in genes involved in neurodevelopment may be associated with altered brain morphology in schizophrenia: preliminary evidence. Psychiatry Res 165(1-2), 1–9.

Gurwitz, David. (2016). Human iPSC-derived neurons and lymphoblastoid cells for personalized medicine research in neuropsychiatric disorders. Dialogues Clin Neurosci 18, 267–276.

Hanson E, Bernier R, Porche K, Jackson FI, Goin-Kochel RP, Snyder LG, Snow AV, Wallace AS, Campe KL, Zhang Y, Chen Q, D’Angelo D, Moreno-De-Luca A, Orr PT, Boomer KB, Evans DW, Kanne S, Berry L, Miller FK, Olson J, Sherr E, Martin CL, Ledbetter DH, Spiro JE, Chung WK; Simons Variation in Individuals Project Consortium. (2015). The cognitive and behavioral phenotype of the 16p11.2 deletion in a clinically ascertained population. Biol Psychiatry 77, 785–93.

Horev G, Ellegood J, Lerch JP, Son YE, Muthuswamy L, Vogel H, Krieger AM, Buja A, Henkelman RM, Wigler M, Mills AA. (2011). Dosage-dependent phenotypes in models of 16p11.2 lesions found in autism. Proc Natl Acad Sci U S A 108, 17076–81.

Iafrate AJ, Feuk L, Rivera MN, Listewnik ML, DonahoePK, Qi Y, Scherer SW, et al. (2004). Detection of large scale variation in the human genome. Nat Genet 36, 949–951.

Jacquemont S, Reymond A, Zufferey F, Harewood L, Walters RG, Kutalik Z, Martinet D, Shen Y, et al. (2011). Mirror extreme BMI phenotypes associated with gene dosage at the chromosome 16p11.2 locus. Nature 478, 97–102.

Jühling F, Kretzmer H, Bernhart SH, Otto C, Stadler PF, Hoffmann S. (2016). metilene: fast and sensitive calling of differentially methylated regions from bisulfite sequencing data. Genome Res 26, 256–62.

Kim D, Pertea G, Trapnell C, Pimentel H, Kelley R, Salzberg SL. (2013). TopHat2: accurate alignment of transcriptomes in the presence of insertions, deletions and gene fusions. Genome Biol 14, R36.

Krueger F, Andrews SR. (2011). Bismark: a flexible aligner and methylation caller for Bisulfite-Seq applications. Bioinformatics 27, 1571–2.

Kumar RA, KaraMohamed S, Sudi J, Conrad DF, Brune C, Badner JA, Gilliam TC, Nowak NJ, Cook EH, Dobyns WB, Christian SL. (2008). Recurrent 16p11.2 microdeletions in autism. Hum Mol Genet 17, 628–638.

Kusenda M, Vacic V, Malhotra D, et al. (2015). The Influence of Microdeletions and Microduplications of 16p11.2 on Global Transcription Profiles. J Child Neurol. 30, 1947–1953.

Langfelder, P. & Horvath, S. (2008). WGCNA: an R package for weighted correlation network analysis. BMC Bioinformatics 9, 559, doi:10.1186/1471-2105-9-559.

Langmead, B. & Salzberg, S. L. (2012). Fast gapped-read alignment with Bowtie 2. Nat Methods 9, 357–359, doi:10.1038/nmeth.1923.

Levy D, Ronemus M, Yamrom B, Lee Y, Leotta A, Kendall J, Marks S, et al. (2011). Rare de novo and transmitted copy-number variation in autistic spectrum disorders. Neuron 70, 886–897.

Li, B. & Dewey, C. N. (2011). RSEM: accurate transcript quantification from RNA-Seq data with or without a reference genome. BMC Bioinformatics 12, 323, doi:10.1186/1471-2105-12-323.

Love MI, Huber W, Anders S. (2014). Moderated estimation of fold change and dispersion for RNA-seq data with DESeq2. Genome Biol 15, 550.

Martin, M. (2011). Cutadapt Removes Adapter Sequences From High-Throughput Sequencing Reads. EMBnet.journal Vol 17, No.1.

Migliavacca E, Golzio C, Männik K, et al. (2015). A Potential Contributory Role for Ciliary Dysfunction in the 16p11.2 600 kb BP4-BP5 Pathology. Am J Hum Genet. 96, 784–796.

Miller DT, Chung W, Nasir R, Shen Y, Steinman KJ, Wu BL, Hanson E. (2015). 16p11.2 Recurrent Microdeletion. Seattle: University of Washington, pp. 1993–2019.

Morrow EM, Yoo SY, Flavell SW, Kim TK, Lin Y, Hill RS, Mukaddes NM, Balkhy S, Gascon G, Hashmi A, Al-Saad S, Ware J, Joseph RM, Greenblatt R, Gleason D, Ertelt JA, Apse KA, Bodell A, Partlow JN, Barry B, Yao H, Markianos K, Ferland RJ, Greenberg ME, Walsh CA. (2008). Identifying autism loci and genes by tracing recent shared ancestry. Science 321, 218–23.

Narayanan B, Soh P, Calhoun VD, Ruaño G, Kocherla M, Windemuth A, Clementz BA, Tamminga CA, Sweeney JA, Keshavan MS, Pearlson GD. (2015). Multivariate genetic determinants of EEG oscillations in schizophrenia and psychotic bipolar disorder from the BSNIP study. Transl Psychiatry 5:e588.

Norata GD, Tavori H, Pirillo A, Fazio S, Catapano AL. (2016). Biology of proprotein convertase subtilisin kexin 9: beyond low-density lipoprotein cholesterol lowering. Cardiovasc Res 112, 429–42.

Poirier S, Prat A, Marcinkiewicz E, Paquin J, Chitramuthu BP, Baranowski D, Cadieux B, Bennett HP, Seidah NG. (2006). Implication of the proprotein convertase NARC-1/PCSK9 in the development of the nervous system. J Neurochem 98, 838–50.

Rashid S, Curtis DE, Garuti R, Anderson NN, Bashmakov Y, Ho YK, Hammer RE, Moon YA, Horton JD. (2005). Decreased plasma cholesterol and hypersensitivity to statins in mice lacking Pcsk9. Proc Natl Acad Sci U S A 102:5374–5379.

Robertson KD. (2005). DNA methylation and human disease. Nature Reviews Genetics. 6, 597–610.

Robinson JT, Thorvaldsdóttir H, Winckler W, Guttman M, Lander ES, Getz G, Mesirov JP. (2011). Integrative genomics viewer. Nat Biotechnol 29, 24–6.

Rosenfeld JA, Coppinger J, Bejjani BA, Girirajan S, Eichler EE, Shaffer LG, Ballif BC. (2010). Speech delays and behavioral problems are the predominant features in individuals with developmental delays and 16p11.2 microdeletions and microduplications. J Neurodev Disord 2, 26–38.

Schulz R, Schlüter KD. (2017). PCSK9 targets important for lipid metabolism. Clin Res Cardiol Suppl 12(Suppl 1), 2–11.

Seidah NG, Mayer G, Zaid A, Rousselet E, Nassoury N, Poirier S, Essalmani R, Prat A. (2008). The activation and physiological functions of the proprotein convertases. Int J Biochem Cell Biol 40, 1111–1125.

Shinawi M, Liu P, Kang SH, Shen J, Belmont JW, Scott DA, Probst FJ, Craigen WJ, Graham BH, Pursley A, Clark G, Lee J, Proud M, Stocco A, Rodriguez DL, Kozel BA, Sparagana S, Roeder ER, McGrew SG, Kurczynski TW, Allison LJ, Amato S, Savage S, Patel A, Stankiewicz P, Beaudet AL, Cheung SW, Lupski JR. (2010). Recurrent reciprocal 16p11.2 rearrangements associated with global developmental delay, behavioural problems, dysmorphism, epilepsy, and abnormal head size. J Med Genet 47, 332–41.

Stark R, Brown G. (2011). DiffBind: differential binding analysis of ChIP-Seq peak data. http://bioconductor.org/packages/release/bioc/vignettes/DiffBind/inst/doc/DiffBind.pdf.

Steinman KJ, Spence SJ, Ramocki MB, Proud MB, Kessler SK, Marco EJ, Green Snyder L, D’Angelo D, Chen Q, Chung WK, Sherr EH; Simons VIP Consortium. (2016). 16p11.2 deletion and duplication: Characterizing neurologic phenotypes in a large clinically ascertained cohort. Am J Med Genet A. 170, 2943–2955.

Sullivan PF, Daly MJ, O’Donovan M. (2012). Genetic architectures of psychiatric disorders: the emerging picture and its implications. Nat Rev Genet 13, 537–51.

Tai DJ, Ragavendran A, Manavalan P, Stortchevoi A, Seabra CM, Erdin S, Collins RL, Blumenthal I, Chen X, Shen Y, Sahin M, Zhang C, Lee C, Gusella JF, Talkowski ME. (2016). Engineering microdeletions and microduplications by targeting segmental duplications with CRISPR. Nat Neurosci 19, 517–22.

Trapnell C, Hendrickson DG, Sauvageau M, Goff L, Rinn JL, Pachter L. (2013). Differential analysis of gene regulation at transcript resolution with RNA-seq. Nat Biotechnol 31, 46–53.

Weber T, Namikawa K, Winter B, Müller-Brown K, Kühn R, Wurst W, Köster RW. (2016). Caspase-mediated apoptosis induction in zebrafish cerebellar Purkinje neurons. Development 143, 4279–4287.

Zhang Y, Pak C, Han Y, Ahlenius H, Zhang Z, Chanda S, Marro S, Patzke C, Acuna C, Covy J, Xu W, Yang N, Danko T, Chen L, Wernig M, Südhof TC. (2013). Rapid single-step induction of functional neurons from human pluripotent stem cells. Neuron 78, 785–98.

Zuffery F, Sherr EH, Beckmann ND, Hanson E, Maillard AM, Hippolyte L, Mace A, Ferrari C, Kutalik Z, Andrieux J, Aylward E, Barker M, Bernier R, Bouquillon S, Conus P, Delobel B, Faucett WA, Goin-Kochel RP, Grant E, Harewood L, Hunter JV, Lebon S, Ledbetter DH, Martin CL, Männik K, Martinet D, Mukherjee P, Ramocki MB, Spence SJ, Steinman KJ, Tjernagel J, Spiro JE, Reymond A, Beckmann JS, Chung WK, Jacquemont S, 16p11.2 European Consortium. (2012). A 600 kb deletion syndrome at 16p11.2 leads to energy imbalance and neuropsychiatric disorders. J Med Genet 49, 660–668.

